# Bcl11b dose-dependently regulates positive selection of CD8 T cells to the virtual memory fate

**DOI:** 10.64898/2026.07.03.731744

**Authors:** Tom Sidwell, Ellen V Rothenberg

## Abstract

Virtual memory T cells are increasingly recognized as a functionally distinct lineage within the CD8 T cell pool, but when and how commitment to the lineage is enforced remain poorly understood. Here we demonstrate that T_VM_ lineage choice is exceptionally sensitive to dosage and repression competence of the key T cell transcription factor Bcl11b. Three different genetic models of slightly reduced Bcl11b each biased CD8 cell development to T_VM_ generation without deregulating effector differentiation. Timed conditional knockouts and adoptive transfers narrowed the developmental window and showed that Bcl11b levels determine diversion to virtual memory fate uniquely during intrathymic positive selection. Whereas total Bcl11b loss disrupts TCR signalling, a <2-fold dose reduction of Bcl11b enhanced selective responses to TCR stimulation. Chromatin accessibility profiling and single cell RNA-seq indicated that Bcl11b dose reduction redirects cells to the T_VM_ fate, from the late cycling fraction of mature CD8SP thymocytes, by a mechanism independent of previously described cytokine-driven pathways.

## Introduction

Virtual memory T (T_VM_) cells are a special lineage of “innate-like” CD8 T cells that have acquired functional and homeostatic features of memory cells without explicit antigen stimulation (Drobek et al., 2018; Haluszczak et al., 2009; Jameson et al., 2015; Kwesi-Maliepaard et al., 2021; Thiele et al., 2020). As the complexity of the CD8 T cell population has become better appreciated, T_VM_ cells have been shown to contribute in important ways as a fast-responding effector subset. They emerge in neonates and in strong cytokine-signaling developmental contexts (Weinreich et al., 2010; Kwesi-Maliepaard et al., 2021; Park et al., 2021), in strongly TCR-signaling intrathymic selection (Drobek et al., 2018; Mahajan et al., 2020), and in a variety of genetic knockout models such as homozygous mutants of *Dot1l* (Kwesi-Maliepaard et al., 2020), *Ezh2* (Dobenecker et al., 2015), *Hdac7* (Kasler et al., 2018), *Dock2* (Mahajan et al., 2020) or *Ikzf3* (Pokhrel et al., 2025). However, there is little insight into the cell-intrinsic steps through which T cells are directed to this choice. Here we introduce a strictly intrathymic path through which developing CD8 T cells are directed to the T_VM_ fate by slightly reduced levels of Bcl11b.

Bcl11b is a zinc finger transcription factor with multiple functions, but in hematopoiesis it is strictly confined to T cells and one subset of innate lymphoid cells (Califano et al., 2015; Walker et al., 2015; Yu et al., 2015). Its expression turns on sharply during the T lineage commitment of early CD4^-^ CD8^-^ (double negative, ‘DN’) thymocytes, before the cells express a T cell receptor (TCR) (L. Li et al., 2010; P. Li et al., 2010), and once expressed it remains active in all subsets of T cells. Working in most if not all mature T cell subsets and during thymic commitment, selection, and CD4/CD8 lineage choice, Bcl11b can bind unique DNA sites but often appears to work primarily as a member of a complex, possibly with GATA3 and very commonly with Runx and ETS factors (Fang et al., 2018; Hosokawa et al., 2018; Kojo et al., 2017; Lorentsen et al., 2018). It associates both with BAF complex components and with NuRD complex components (Cismasiu et al., 2005; Kadoch et al., 2013). However, the genes regulated by Bcl11b appear quite different in different T-cell contexts. Although some functional target genes are shared, the impact of Bcl11b loss appears to be very cell-type and stage-dependent (reviewed in (Sidwell and Rothenberg, 2021)).

Despite its ubiquity in T cells, the amplitudes of Bcl11b expression are strictly set at different characteristic levels for T cells in different subsets and different stages, appearing to decrease during T-cell activation (Dubuissez et al., 2016; Zhang et al., 2012). Recent evidence suggests that changes in human T and NK cells could result from altered Bcl11b levels (Holmes et al., 2021; Kumar et al., 2025; Liao et al., 2023). However, virtually all actions inferred for Bcl11b have been identified via experimental strategies where it was homozygously knocked out or functionally disrupted. Only a few hypomorphs have been characterized experimentally (Hirose et al., 2015; Kojo et al., 2018; Matsumoto et al., 2024).

Varying transcription factor dose has repeatedly been shown to have the capacity to alter hematopoietic outcomes through graded gene expression effects (Allman et al., 2006; Heshusius et al., 2022; Hohaus et al., 1995; Mak et al., 2011; Peters et al., 2023; Wang et al., 1996). In accord, mutations in humans causing contextual and quantitative impacts upon gene expression rather than total loss of function can be highly significant for human disease (Loos, 2020; Watanabe et al., 2019). In mice, T cell development has been shown to be affected significantly by changes of 2x or less from the normal level of GATA3 (Scripture-Adams et al., 2014; Xu et al., 2013), Runx1 (Shin et al., 2023), or ETS1 (Chandra et al., 2023). Here, we show that there is a critical stage of intrathymic development at which a modest reduction of Bcl11b permanently alters the fates of developing CD8 T cells and shunts them to fates as pre-poised naïve and virtual memory cells.

## Results

### Bcl11b dose reduction expands the CD8 T cell central memory compartment

Results of previous studies suggested that Bcl11b might have a dose-dependent role in T cells. First, in comparing RNA expression in *Bcl11b* homozygous knockout thymocytes to their normal counterparts, the heterozygous DN3 cells also differed markedly from Bcl11b-replete controls, although this was dwarfed by the effect of the homozygous knockout (Hosokawa et al., 2018) (**Figure 1 – figure supplement 1a, Supplementary table 1**). Second, in a genetic model where about 10-20% of cells express *Bcl11b* monoallelically, the mature fates of the monoallelically expressing cells appeared to be skewed relative to the biallelically expressing cells in the same mice (Ng et al., 2018) (see below). We therefore sought to assess whether modestly reduced Bcl11b dose truly alters fate choice within the T cell lineages. Three genetic strategies were used, as summarized in **Figure 1 – figure supplement 2a,b** (with the three primary models labelled), a germline heterozygote of *Bcl11b* gene disruption, a homozygote of a *Bcl11b* enhancer region deletion, and a conditional heterozygous knockout of *Bcl11b* starting from the CD4^+^CD8^+^ (DP) stage in the thymus.

### Germline *Bcl11b* heterozygotes

We first utilised a germline Bcl11b dosage reduction mouse model. One allele of the *Bcl11b* gene in these animals is disrupted by insertion of mCherry in place of the first exon (Rothenberg et al., 2016). This allele is functionally Bcl11b-null and causes lethality if homogyzous (**Figure 1 – figure supplement 2a, b “Germline deficiency”**), but the *Bcl11b* regulatory elements remain intact (**Figure 1 – figure supplement 2c**). Thus, the mCherry reporter in this allele tracks locus transcription, and all heterozygous T cells beyond the DN2 intrathymic stage fluoresce red. When fetal hematopoietic precursors homozygous for the allele are allowed to develop in chimeras, the mCherry-expressing cells switch to an NK-like fate (**Figure 1 – figure supplement 1b**), like any other homozygous *Bcl11b* knockout cells (P. Li et al., 2010; Liao et al., 2023). The mCherry-disrupted *Bcl11b* heterozygotes will be referred to as *Bcl11b^+/-^*.

Bcl11b heterozygosity had no observed impact on overall cellularity in the spleen, while total CD8 T cell numbers were modestly reduced relative to littermate controls (**Figure 1a**). However, within the splenic CD8 T cell compartment, there was a marked population shift from controls. There was an approximately 3-fold increase in the frequency of CD44^+^CD62L^+^ central memory T (T_CM_)-phenotype cells in heterozygous mice relative to controls, at the expense of naïve (T_N_) cells (**Figure 1b**). Further, even the cells remaining in the *Bcl11b^+/-^* “naive” CD8 T cell gate showed elevated CD44 expression relative to controls (**Figure 1b**, **Figure 1 – figure supplement 3a**). In contrast, the percentage of CD44^+^ CD62L^-^ effector-memory cells was less consistently affected (**Figure 1b**). Thus, Bcl11b is haploinsufficient for control of the ratio of T_N_ : T_CM_ phenotype cells in the CD8 T cell pool.

**Figure 1.**
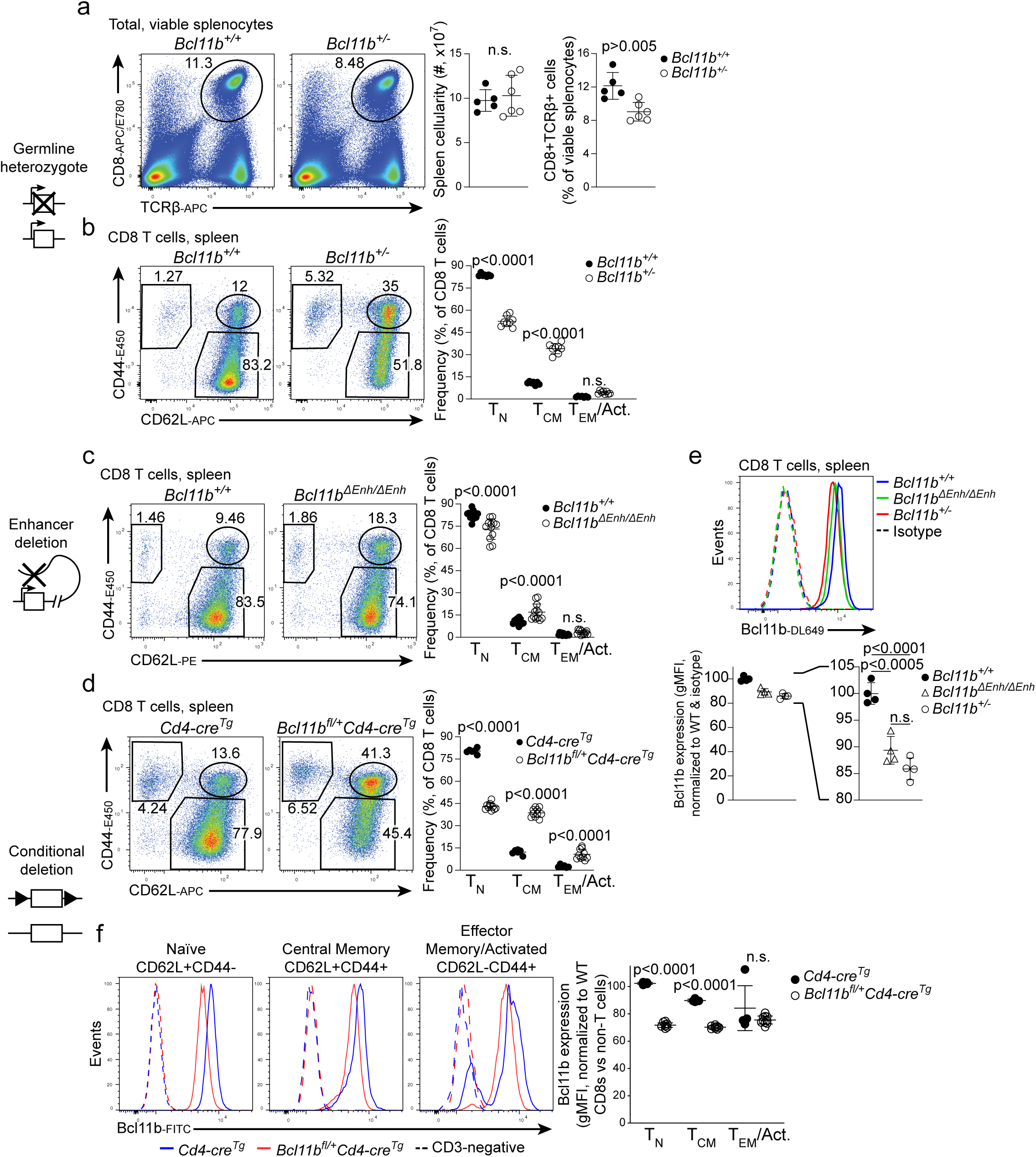
Bcl11b dose restrains the CD8 T cell memory:naïve ratio. **a –b.** Flow cytometric characterisation of Bcl11b haploinsufficient and wildtype control splenic CD8 T cells. **a.** Representative flow cytometry plots of CD8 T cell frequencies (left) and quantification of total splenic cellularity and CD8 T cell frequencies (right). **b.** Representative flow cytometry plots of naïve (CD62L+ CD44-low, T_N_), central memory (CD62L+CD44+, T_CM_) and activated/effector memory (CD62L- CD44+, T_EM_/Act.) CD8 T cells (left), quantified right. **c.**, as for b., comparing *Bcl11b^ΔEnh/ΔEnh^* enhancer mutant and control splenic CD8 T cells. **d.** As for b. and c., comparing *Bcl11b^fl/+^ Cd4-cre^Tg^* conditional heterozygote and *Cd4-cre^Tg^* control splenic CD8 T cells. **e.** Comparison of Bcl11b expression between co-stained Bcl11b haploinsufficient, enhancer mutant and wildtype control splenic CD8 T cells. Upper, histograms of expression from representative samples, lower, quantification. **f.** Left, representative histograms of Bcl11b expression by conditionally haploinsufficient and Cre-expressing control CD8 T cells of naïve, T_CM_ and activated/T_CM_ phenotypes, quantified right. Significance tested using Student’s two-tailed t test (a) or one-way ANOVA with Šidák’s (b-d, f) or Tukey’s (e) correction for multiple comparisons.

The CD4-positive T cell lineages from these Bcl11b-haploinsufficient spleens appeared less affected than CD8 T cells. Both Foxp3^-^ conventional and Foxp3^+^ regulatory CD4 T (T_REG_) cells were indistinguishable in cell numbers between *Bcl11b^+/-^* and littermate *Bcl11b^+/+^* control spleens (**Figure 1 – figure supplement 3b, c**). However, *Bcl11b*^+/-^ T_REG_ cells displayed a modest decrease in frequency of naïve phenotype cells compared with controls (**Figure 1 – figure supplement 3c**). Both the CD62L^+^CD44^low^ naïve CD4conv and the resting (CD62L^+^KLRG1^-^) T_REG_ cell populations in heterozygotes showed a trend toward higher CD44 expression than in controls (**Figure 1 – figure supplement 3b, c**). Thus, the impact of Bcl11b haploinsufficiency on T cell activation status in homeostasis is not unique to the CD8 T cell lineage, though it presents most dramatically there.

### Alternative model of Bcl11b dosage reduction: enhancer deletion

To rule out the possibility that the mCherry-disrupted *Bcl11b* allele was causing these abnormalities through an aberrant gene product, we examined an alternative model of reduced Bcl11b expression, an enhancer mutant (*Bcl11b^ΔEnh^*) in which the gene body remains completely intact (**Figure 1 – figure supplement 2b**). The “major peak” enhancer of *Bcl11b* (Li et al., 2013), ∼800kb downstream of the gene body, is required for the timely initiation of Bcl11b expression within the thymic double negative compartment (Ng et al., 2018; Pease et al., 2024). In *Bcl11b^ΔEnh^*^/+^ heterozygotes, we previously found that a minority of cells never activate expression from the enhancer-mutant allele,yielding a permanent minority of monoallelically expressing T cells (Ng et al., 2018). Those monoallelically expressing cells were underrepresented in the T_REG_ cell compartment and overrepresented among γδ T cells and CD8^+^ T cells with a putatively antigen-experienced CD44^+^ phenotype (Ng et al., 2018). When we bred the *Bcl11b^ΔEnh^* allele to homozygosity, both copies of the enhancer-deleted allele were likely to be activated later than the enhancer-replete allele, but all cells that developed successfully into T cells ultimately expressed some *Bcl11b*. As shown below, though, loss of this specific enhancer element caused cells to express a slightly lower level of Bcl11b.

Homozygous *Bcl11b* enhancer mutant mice had normal frequencies of major CD4/CD8 thymocyte subsets relative to controls (**Figure 1 – figure supplement 3d**), and in the spleen, cell numbers and CD8 T cell frequencies were unaffected by loss of the distal *Bcl11b* enhancer region (**Figure 1 – figure supplement 3e**). However, within the CD8 T cell compartment, these *Bcl11b^ΔEnh/ΔEnh^* mice slightly but reproducibly displayed increased numbers and frequencies of T_CM_-phenotype cells compared with controls (**Figure 1c**), with reduction of the T_N_ populations.

### Conditional knockout model of Bcl11b dosage reduction

Both the germline *Bcl11b^+/-^* and the germline *Bcl11b^ΔEnh/ΔEnh^* dosage reduction models should differ from wildtype in Bcl11b levels starting from the T cell lineage commitment stage in DN2 thymocytes, when *Bcl11b* expression is first activated (see **Figure 1 – figure supplement 2a**, timing when mutations take effect). Indeed, the *Bcl11b^ΔEnh^* allele can alter DN thymocyte biology (Pease et al., 2024). To evaluate whether the Bcl11b dosage reduction phenotype depended on effects starting before the DP stage, we tested a conditional knockout model, in which transgenic CD4-Cre (Lee et al., 2001) was used to excise one copy of *Bcl11b* exon 4 in *Bcl11b^fl/+^* heterozygous cells, from the thymic pre-DP (DN4) to DP stage (Shi and Petrie, 2012). The number and frequency of the major thymic CD4/CD8 subsets appeared equivalent between *Bcl11b^fl/+^ Cd4-cre^Tg^* and Cre-control animals (**Figure 1 – figure supplement 3f**), and total splenic cellularity appeared normal, although CD8 T cells were slightly reduced (**Figure 1 – figure supplement 3g**) with lower levels of CD8α and CD3ε expression per cell than controls (**Figure 1 – figure supplement 3h**). Again, however, the CD8 cells in this dose-reduction model were shifted to a T_CM_-like phenotype at the expense of cells with a naïve-like phenotype **(Figure 1d),** with increased CD44 expression on the cells remaining within the “naïve” gate relative to those in Cre-expressing controls. Thus, all three models of *Bcl11b* dosage reduction produced a shift from naïve to memory-phenotype CD8 cells, and no effects of Bcl11b reduction in the early DN stages of T cell development were required for this effect.

### Less than twofold differences in Bcl11b expression alter CD8 T cell development

We used intracellular staining to compare Bcl11b expression levels between T cells of these three dosage-reduced Bcl11b genotypes. Heterozygosity might be expected to reduce Bcl11b levels by twofold, but the differences in Bcl11b expression were reproducibly smaller than this. They were therefore measured under highly controlled conditions for accuracy (see **Methods**). In splenic CD8 T cells, Bcl11b protein levels clearly resolved with levels in wildtype > enhancer mutant > heterozygous knockout cells overall (**Figure 1e**). Thus, the proportional frequency of T_CM_-like CD44^+^ CD62L^+^ cells among the CD8 compartment, heterozygous knockout > enhancer mutant > wildtype, was inversely correlated with Bcl11b protein levels. However, for both *Bcl11b^+/-^* and *Bcl11b^fl/+^ Cd4-cre^Tg^* models (**Figure 1e, f**), the heterozygotes still expressed over 70% as much Bcl11b protein as *Bcl11b^+/+^* controls, possibly because Bcl11b slightly represses expression of its own coding locus (Hosokawa et al., 2018). As expected, Bcl11b protein levels in controls were highest in naïve CD8 cells and reduced in T_CM_ and T_EM_ gated cells. While Bcl11b levels were comparably low in conditional heterozygote and control T_EM_ cells (**Figure 1f**), per subset, heterozygotes showed 70% and 80% of control levels of Bcl11b in naïve and central memory T cells, respectively (**Figure 1 – figure supplement 3i**). These results suggest that, *in vivo,* CD8 T cell subset choice can be altered by a very small reduction in Bcl11b.

### Normal dose-dependent function of Bcl11b in CD8 T cells depends on integrity of N-terminal corepressor-recruitment domain

Bcl11b binds to three kinds of targets when it functions: genomic DNA at specific sites (Avram et al., 2002; Fang et al., 2018), other sequence specific transcription factors (such as Runx1 (Hosokawa et al., 2018) and the Nr2f “COUP-TF” family (Avram et al., 2000)), and chromatin modifying proteins (see Introduction; rev. in (Sidwell and Rothenberg, 2021)). To test whether the exceptionally dosage sensitive function of Bcl11b depended on recruitment of corepressor complexes as well as on DNA binding, we took advantage of a recently described Bcl11b point mutation model, Bcl11b-R3S (Goos et al., 2019)(**Figure 1 – figure supplement 2a, b**). This patient-derived mutation disrupts the N-terminal FOG repressor domain that Bcl11b uses to recruit the NuRD and Polycomb complexes, but without altering the DNA binding zinc fingers (Cismasiu et al., 2005; Goos et al., 2019). As the homozygous allele results in perinatal lethality, E17 fetal thymocytes were analyzed (**Figure 2a-e**). Surprisingly, development of DN thymocytes all through the DN3 to DN4 transition was close to normal (**Figure 2a-c**), although generation of DP thymocytes was inhibited or delayed in *Bcl11b^R3S/R3S^* homozygotes (**Figure 2d**). However, we confirmed that the R3S mutation indeed interfered with repression of specific Bcl11b targets, as fetal DN thymocytes from *Bcl11b^R3S/R3S^* homozygotes failed to repress expression of an early Bcl11b silencing target, c-Kit, in a timely manner (**Figure 2e**).

**Figure 2.**
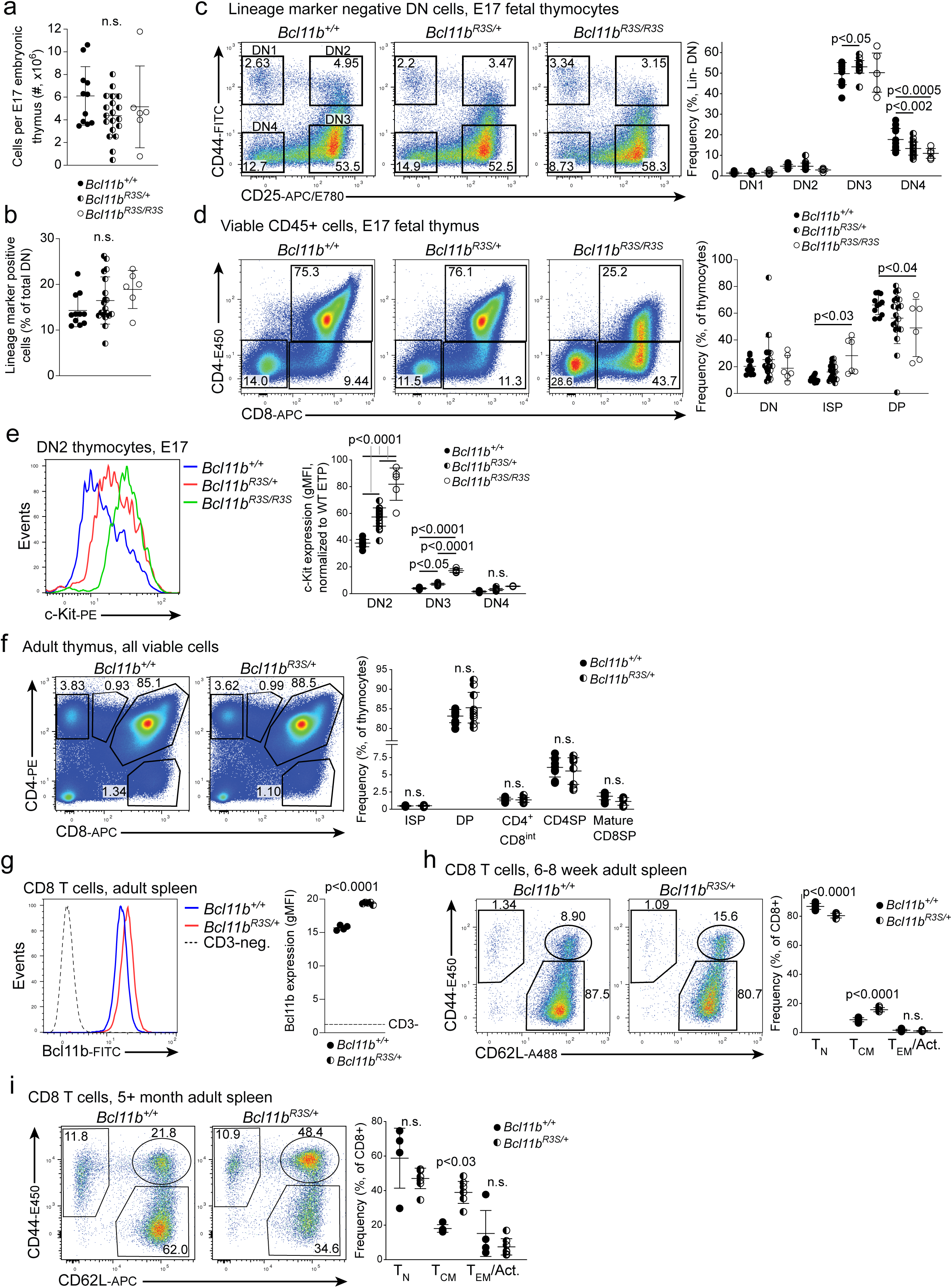
**a.-e.** E17 fetal thymuses from *Bcl11b^+/+^*, *Bcl11b^R3S/+^* and *Bcl11b^R3S/R3S^* embryos analyzed by flow cytometry. **a.** Total thymic cellularity. **b.** Frequency of lineage marker positive cells. **c.** Representative flow cytometry plots (left) and frequency (right) of DN subsets. **d.** Representative flow cytometry plots (left) and frequency (right) of CD4/CD8 expressing populations. **e.** Representative histograms of DN2 cells (left), and quantification among DN2-DN4 cells (right) of c-Kit expression measured by flow cytometry. **f.** Representative flow cytometry plots (left) and quantification (right) of major CD4/CD8 expressing populations among adult *Bcl11b^R3S/+^* and littermate control thymuses. **g.** Bcl11b quantification by flow cytometry among splenic CD8 T cells. **h.-i.** T_N_, T_CM_ and T_EM_ frequencies among splenic CD8 T cell populations, **h.** 6-8 week old and **i.** 5-8 month old mice. Significance tested using one-way ANOVA with Tukey’s (a- e) or Šidák’s (f, h, i) correction for multiple comparisons, or Student’s two-tailed t test (g).

Heterozygous *Bcl11b^R3S/+^* mice were viable, and the frequencies of the major CD4/CD8 expressing populations did not significantly differ from wildtype littermate control thymuses (**Figure 2f**). The overall Bcl11b protein content of *Bcl11b^R3S/+^* CD8 T cells was about 30% higher than that of control cells (**Figure 2g**), in clear contrast to the three dosage reduction models just described. Nevertheless, in the heterozygous R3S mutant animals, the T_CM_-phenotype population within the splenic CD8 compartment was expanded compared with wildtype controls, again with increased CD44 expression in the naïve subset (**Figure 2h**), resembling a weaker form of the dose-reduced models. This effect, visible at 6-8 wk old, became quite strong in older animals (**Figure 2i**). Thus, even when the total level of Bcl11b protein was not reduced, a reduced ability of the Bcl11b pool to recruit corepressor was sufficient to bias CD8 cells to central memory-like phenotype, mimicking Bcl11b dosage reduction.

### Bcl11b haploinsufficient CD8 T cells display multiple central or virtual memory features and functions

To verify whether the expanded CD44^+^ CD62L^+^ subset in Bcl11b-reduced CD8 T cell populations were truly memory-like, we assessed them for multiple additional functional and gene-expression features that discriminate between naïve and memory T cells in wildtype mice. The expression of CD127 and KLRG1 was comparable between *Bcl11b*-haploinsufficient naïve, central memory and effector memory CD8 T cells and the corresponding wildtype controls (**Figure 3a**). Similarly, levels of Eomes as well as TCF1 – transcription factors essential for the establishment and maintenance of CD8 T cell memory (Banerjee et al., 2010; Jeannet et al., 2010) – were as high in Bcl11b haploinsufficient CD44^+^ CD62L^+^ CD8 T cells as in T_CM_-phenotype cells from wildtype control mice (**Figure 3b**). The expanded memory-like cells also showed functional competence like wildtype memory cells, based on cytokine production after PMA/ionomycin stimulation between *Bcl11b^+/-^* and control CD8 T cells. Comparing populations with the same naïve/memory marker status, the frequencies of cells able to express these cytokines upon short-term stimulation were equally low in naïve subsets and equally high in memory subsets from Bcl11b haploinsufficient and wildtype control mice alike (**Figure 3 – figure supplement 1a**). Thus, because of their increased percentages of cells with T_CM_ phenotype, the total CD8 T cell compartments from Bcl11b haploinsufficient mice contained much higher frequencies of cells producing IFNγ and/or TNFα on short-term stimulation than wildtype controls (**Figure 3c**).

**Figure 3.**
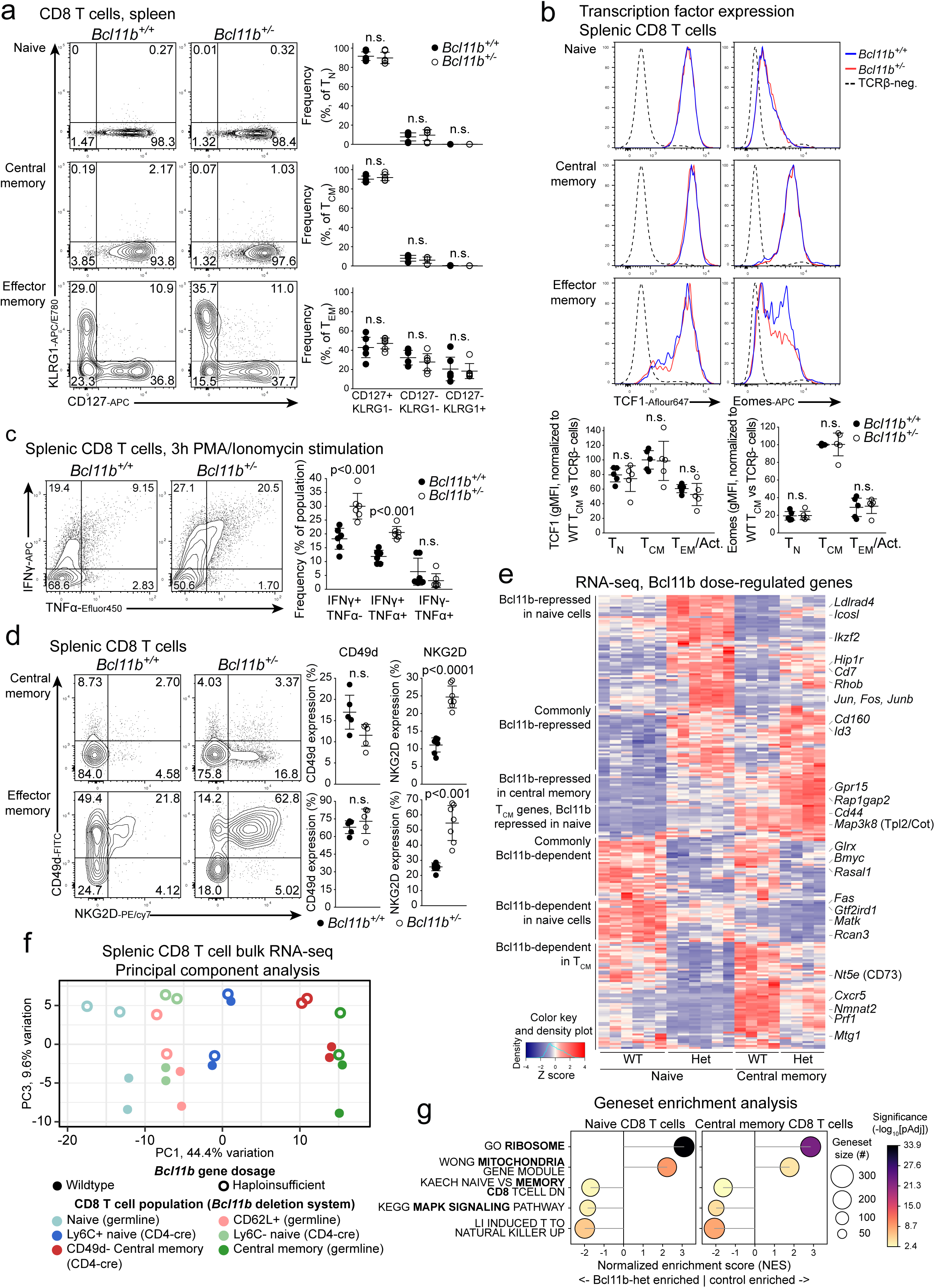
Bcl11b haploinsufficiency expands the virtual memory compartment. **a.-d.** Flow cytometric characterisation of the splenic CD8 T cell compartments of *Bcl11b^+/-^* and littermate control mice. **a.** Left, representative plots and right, quantification, of KLRG1 and CD127 (IL-7 receptor alpha chain) expression by T_N_, T_CM_ and T_EM_ CD8+ subsets. **b.** Upper, representative histograms and lower, quantification, comparing TCF1 and Eomes expression by T_N_, T_CM_ and T_EM_ CD8 T cells of each genotype. **c.** Left, representative plots and right, quantification of IFN-γ and TNF-α expression following stimulation with PMA and ionomycin with brefeldin. **d.** Left, representative flow cytometry plots and right quantification of expression of virtual memory T cell identity-associated proteins CD49d and NKG2D by T_CM_ and T_EM_ CD8 populations. **e.-g.** Bulk RNA-seq of splenic CD8 T cell populations in germline and conditional Bcl11b haploinsufficiency models. **e.** Z-score plot of all genes significantly differentially expressed between genotypes in naïve or central memory phenotype cells, select genes labelled. **f.** Principal component analysis of sequenced samples using 10% most variant genes. **g.** Summary of geneset enrichment analysis (GSEA) of select pathways significantly enriched between genotypes in naïve and T_CM_ phenotype samples. Significance tested using one-way ANOVA with Šidák’s correction (a-c), or Student’s two-tailed t test (d).

In wildtype, specific pathogen-free mice, a large proportion of T_CM_-phenotype cells are not genuinely antigen-experienced, and so are termed ‘virtual’ memory T (T_VM_) cells (Haluszczak et al., 2009; Thiele et al., 2020). Such T_VM_ cells lack CD49d, which denotes ‘true’ (activation-induced) memory cells, but they tend to express NKG2D, a protein expressed by some T_VM_ cells and effector memory (T_EM_) cells. Using these markers to assess the Bcl11b-haploinsufficient expanded subset, we determined that there was no expansion of true CD49d^+^ memory cells in Bcl11b haploinsufficient CD8 T cells (**Figure 3d**). Cells with a T_CM_ phenotype in *Bcl11b^+/-^* mice displayed increased frequencies of NKG2D expression compared with *Bcl11b^+/+^* controls, but no increase in CD49d_;_ thus, most could be antigen-inexperienced (**Figure 3d**). Likewise, in CD8 T cells from *Bcl11b^fl/+^ Cd4-cre^Tg^*mice, the number of CD49d+ ‘true’ memory cells was not increased (**Figure 3 – figure supplement 1b**). Thus, the expanded T_CM_-like populations of *Bcl11b* haploinsufficient CD8 T cells consisted of T_VM_ cells.

Bcl11b haploinsufficiency also affected the CD8 cells remaining naive, i.e., cells with high TCF1, low Eomes, and low ability to express IFNγ and TNFα. Not only did these cells show up-shifted CD44 expression compared to wildtype controls, but they also had a strongly increased percentage of cells expressing Ly6C (**Figure 3 – figure supplement 1c**). In addition to memory phenotype cells, Ly6C is expressed by a discrete subset of naïve peripheral T cells that is generated through high TCR and type I interferon signaling within the thymus (Jergović et al., 2021; Ju et al., 2021). Thus, Bcl11b dosage reduction affected naïve subsets themselves as well as expanding the population of T_VM_ cells in the CD8 compartment overall.

### Bcl11b dose reduction generates both naïve and T_VM_ cells with altered transcriptomes

Although they appeared broadly similar to control populations in phenotype and function, reduced Bcl11b dosage did affect gene expression in both naïve and T_VM_ subsets, as shown by bulk RNA-seq of sorted naïve (CD44^-^CD62L^+^), T_CM/VM_ (CD44^+^CD62L^+^) and pooled total CD62L^+^ splenic CD8 T cells from *Bcl11b^+/-^* or *Bcl11b^fl/+^Cd4-cre^Tg^* and littermate or Cre-expressing control mice, respectively (**Figure 3e-g**). Effects on gene expression in each subset were mild but reproducible (genes differentially expressed by genotype shown in **Figure 3e**). Principal component (PC) analysis of expression of the 10% of genes most variable between samples showed that the first PC resolved the samples according to naïve/memory status and the third PC resolved their genotypes. There was excellent agreement between the two models of *Bcl11b* dosage reduction and, especially in the CD4-Cre model, both naïve and central memory haploinsufficient cells clearly separated from the controls (**Figure 3f**). Gene set enrichment analysis (Subramanian et al., 2005) (GSEA), using Molecular Signatures Database (MSigDB) gene sets from the immunological signature, curated or hallmark gene set collections, confirmed that multi-gene signatures of memory T cell identity, induced T-to-NK conversion, and MAP kinase signaling were significantly enriched in *Bcl11b* haploinsufficient CD8 T cells relative to controls (**Figure 3g**). While ribosomal and mitochondrial function gene sets were strikingly down in heterozygotes (**Figure 3g**; i.e. control-enriched), individual genes in these sets did not reach significance thresholds (**Supplementary tables 2, 3**). Notably, though, signatures of signals known to expand T_VM_ cells in other models, such as IL-4, IL-15 and interferon signaling, were not significantly enriched in *Bcl11b* heterozygous cells, suggesting a distinct pathway.

In CD8 cells with a CD62L^+^ CD44^low^ “naïve”-like phenotype, 120 genes were detectably differentially expressed (pad j< 0.05 and fold change > 1.5 up or down, i.e. |log2 FC| > 0.584), with 64 genes higher in the Heterozygotes, and 51 reduced in expression relative to controls (**Supplementary table 2**). In the T_CM/VM_ compartment of CD8 T cells, 38 upregulated and 32 downregulated genes distinguished the *Bcl11b* heterozygotes from controls (**Supplementary table 3**). Bcl11b dose reduction affected many genes in subset-specific ways (**Figure 3e**). However, expression of genesets associated with T to NK transformation of homozygously *Bcl11b*-deleted T cells (‘ITNK’ signature) (P. Li et al., 2010) and MAPK and T cell receptor (TCR) signaling (**Figure 3g**) were enriched in both naïve and T_VM_ subsets of *Bcl11b* heterozygous CD8 cells.

Interestingly, GSEA showed (**Figure 3 – figure supplement 1d**) that the genes differentially expressed in Bcl11b haploinsufficient CD8 T cells included some genes also impacted by homozygous and heterozygous Bcl11b loss in DN3 thymocytes (Hosokawa et al., 2018) (described **Figure 1 – figure supplement 1a**) or homozygous loss of Bcl11b among mature CD8 T cells (Helm et al., 2023). Although only a few genes were involved in this overlap, some conserved Bcl11b repression targets were evident (**Figure 3e**). *Cd7* and *Osbpl5* were significantly Bcl11b repressed in all conditions, and *Rhob*, *Hip1r*, *Cd9, Cd160* and *Cd44* were also implied to be Bcl11b repressed across at least three contexts. Thus, Bcl11b haploinsufficient mature CD8 T cells shared some signature gene expression features of Bcl11b-null cells across T cell development.

### Bcl11b dose reduction increases TCR responsiveness and speeds proliferation of mature naïve T cells

An expanded T_VM_ compartment in homeostasis might result from increased TCR signal sensitivity in naïve T cells, to augment the activation response to self-antigens, or from increased proliferation in response to cytokine or T-cell receptor (TCR) signals. To compare the sensitivities to TCR signaling of naïve Bcl11b dose-reduced T cells and naïve wildtype controls, we stimulated them with different doses of anti-CD3 antibody, assessing expression of known TCR-responsive molecules after 16 hr of stimulation. The *Bcl11b^+/-^* naïve CD8 cells were markedly more sensitive than wildtype T cells, requiring lower doses of anti-CD3 to activate CD44, CD69 and CD25 than the controls (**Figure 3 – figure supplement 1e**). We considered the possibility that this might be due to positive Bcl11b dependence of CD5 (Hosokawa et al., 2018), a TCR-upregulated cell surface receptor that can dampen TCR sensitivity through a negative feedback (Azzam et al., 1998). However, CD5 upregulation was unaffected by Bcl11b dose in these peripheral CD8 T cells. Thus, reduced levels of Bcl11b must enhance TCR signaling sensitivity of naïve CD8 cells by another mechanism.

Naïve CD8 T cells from *Bcl11b^+/-^* mice also displayed a proliferative advantage compared with controls when responding to anti-CD3. At day 3 of culture, Bcl11b heterozygous CD8 and CD4 naïve T cells each underwent slightly faster proliferation, measured by cell number and dilution of division tracing dye (**Figure 3 – figure supplement 1f, g**). Faster proliferation was also seen when naïve CD8 T cells from *Bcl11b^+/-^* and littermate control mice were compared in cultures with IL-7 alone, absent exogenous TCR stimulation. Again, more *Bcl11b^+/-^* cells entered into division, and these clones underwent more divisions, than did *Bcl11b^+/+^* controls (**Figure 3 – figure supplement 1h**). Thus, normal levels of Bcl11b restrain proliferation of naïve CD8 and CD4 T cells in response to TCR stimulation and cytokines alike.

### Bcl11b dosage effect is cell-intrinsic and does not require lymphopenia

To verify whether the shift from naïve to virtual memory bias in the Bcl11b haploinsufficient T cells is caused by a cell-intrinsic mechanism, we carried out developmental chimera experiments transferring *Bcl11b^+/+^* or *Bcl11b^+/-^* bone marrow precursor cells into busulfan-conditioned wildtype recipients, allowing *Bcl11b^+/+^* and *Bcl11b^+/-^* donor cells to develop in parallel in the same mature T-replete environment. We generated the bone marrow chimeras under competitive conditions, exploiting the diverse Bcl11b fluorescent reporter strains we developed previously as controls (Ng et al., 2018; Pease et al., 2024). Thus, Bcl11b-wildtype BM competitors with mCitrine as nondisruptive reporter were mixed with test BM cells from either Bcl11b^+/-^ (with mCherry disrupting Bcl11b), or Bcl11b-wildtype control (with mCherry as nondisruptive reporter) genotypes, and the mixtures used to repopulate busulfan-conditioned wildtype hosts (**Figure 4a**). At 8 weeks post reconstitution, the *Bcl11b^+/+^* test cells and the wildtype competitors in all chimeras all showed the same balance of preponderantly naïve CD8 T cells with a minority of T_VM_/T_CM_ cells (**Figure 4a, lower** plot), whereas the Bcl11b-heterozygous disruptive mCherry^+^ test cells in the mixed chimeras alone showed the shift to T_VM_ generation (**Figure 4a, upper** plot).

**Figure 4.**
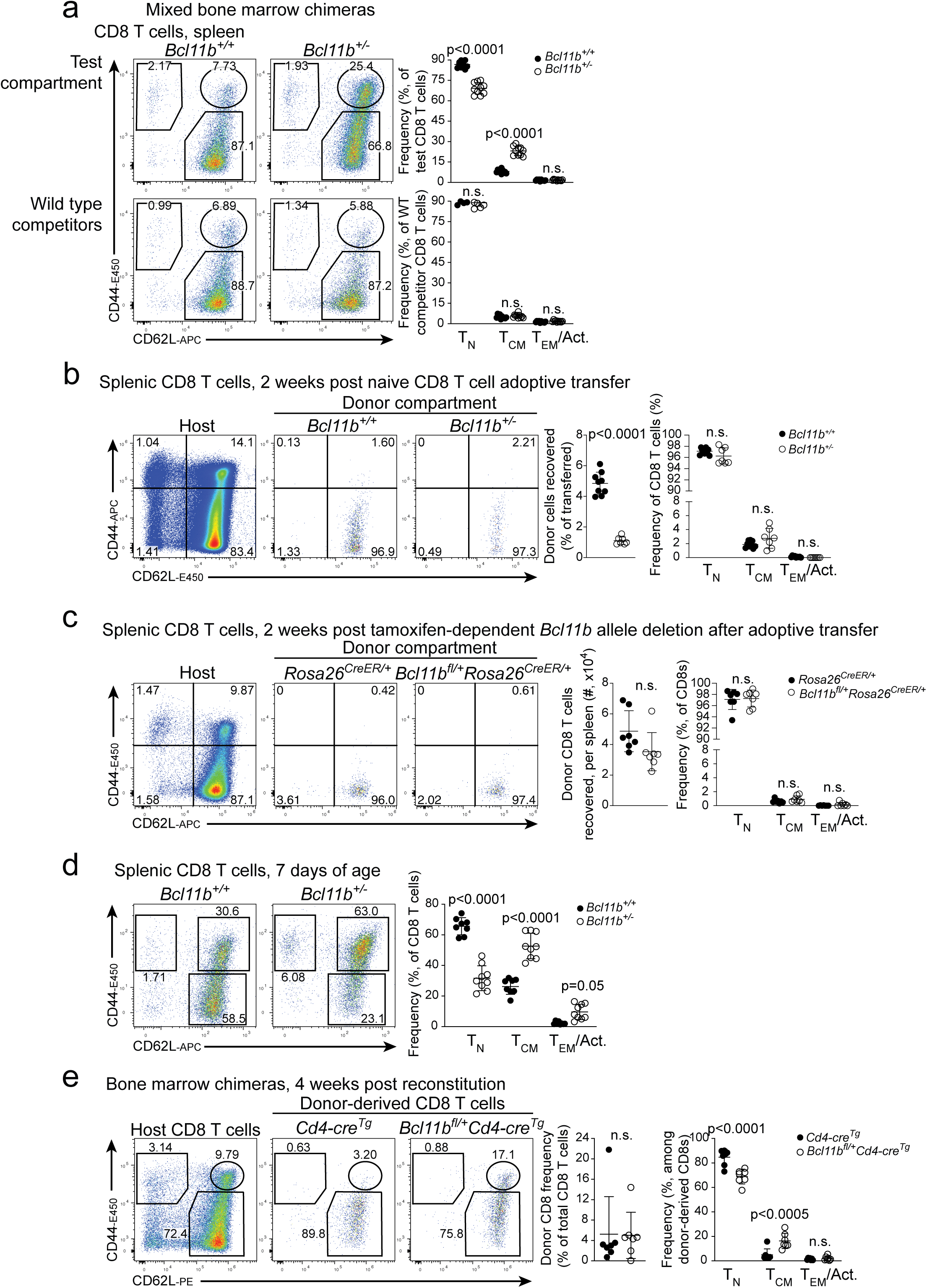
Bcl11b dose determines T_VM_ compartment size before T cells enter the periphery. The T_VM_ generating potential of wildtype and *Bcl11b*-haploinsufficient cells were determined in transfers of the indicated cell populations to wildtype hosts and in the postnatal ontogenic timecourse of T_VM_ production in mice with different *Bcl11b* genotypes. No recipient animals were depleted of pre-existing host CD8 T cells for these experiments. **a.** Representative flow cytometry plots of splenic CD8 T cells compartments of mixed bone marrow chimeras (left). Upper, from *Bcl11b^+/-^* or control donors, or lower, wildtype competitors. Right, quantification of T_N_, T_CM_ and T_EM_ frequencies among test compartments (upper) vs their wildtype competitors (lower). **b.** Naïve CD8 T cells from *Bcl11b^+/-^* or wild type control spleens were adoptively transferred into wild type hosts, and donor cells analysed two weeks later. Left, representative flow cytometry plots of host and donor CD8 T cells. Right, quantification of recovered cells and the frequency of T_N_, T_CM_ and T_EM_ phenotypes. **c.** 400,000 naïve CD8 T cells from *Bcl11b^fl/+^Rosa26^CreERT2/+^* or *Rosa26^CreERT2/+^* control mice were transferred into new hosts, which were subsequently administered tamoxifen. Right, representative flow cytometry plots of host and donor CD8 T cell phenotypes two weeks post transfer. Right, quantification of recovered cells and T_N_, T_CM_ and T_EM_ frequencies. **d.** Splenocytes from 7 day old *Bcl11b^+/-^* and littermate control pups were analysed by flow cytometry. left, representative flow plots and, right, quantification of T_N_, T_CM_ and T_EM_ cell frequencies among CD8 T cells. **e.** Bone marrow chimeras with either *Bcl11b^fl/+^Cd4-cre^Tg^* or Cre-control donor cells analysed four weeks post reconstitution. Left, representative flow plots of host and donor CD8 T cell phenotypes. Right, quantification of donor cell frequency among total (host + donor) CD8 T cells, and of of T_N_, T_CM_ and T_EM_ cell frequencies among donor cells by genotype. Significance tested using one-way ANOVA with Šidák’s correction (a-e), or Student’s two-tailed t test (b, c, e).

We conditioned the hosts with busulfan rather than whole body irradiation because it does not induce an inflammatory environment and spares host mature T cells (Hsieh et al., 2007; Montecino-Rodriguez et al., 2019; Yeager et al., 1991). Thus, the propensity of Bcl11b haploinsufficient cells toward T_VM_ differentiation occurred absent the lymphopenia which could otherwise drive homeostatic proliferation-linked conversion to memory-phenotype T cells. However, the absolute frequency of donor T_VM_ cells in chimeras was not as high as in the endogenous populations of mice of the same genotypes in steady state (cf. **Figure 2b, d**), possibly because the bone marrow chimeric model does not recapitulate the highly T_VM_-genic neonatal stage (Smith et al., 2018), and/or due to competition with host memory cells for access to the niche. Thus, Bcl11b dosage reduction causes a shift to T_VM_ production, without a requirement for a lymphopenic environment, and via a cell-autonomous mechanism.

### Bcl11b haploinsufficient cells do not convert more readily from naïve to virtual memory in the periphery

As the “naïve” phenotype cells in the Bcl11b dosage-reduced mice appeared sensitized for activation and growth, we hypothesized that they might be continuously converting to T_VM_ in the periphery, possibly through increased response to contact with higher affinity self antigens (Drobek et al., 2018; Mahajan et al., 2020). Alternatively, virtual memory T cells have been reported to emerge from a possible intrathymic developmental branch point (Miller et al., 2020). T_VM_ cells that are generated by signals in the periphery have been reported to express CCR2 (Hou et al., 2021). However, *Ccr2* expression was low in all T_CM/VM_ samples, regardless of genotype (**Supplementary table 3**), suggesting there is no specific expansion of peripherally derived T_VM_ cells.

To test this directly, we isolated naïve mature CD8 T cells from spleens and lymph nodes of *Bcl11b^+/-^* or wildtype Bcl11b-mCherry reporter control mice and adoptively transferred these cells into wildtype hosts. Two weeks later, we assessed the phenotype of donor-origin cells. There was a distinct defect in the persistence of *Bcl11b^+/-^* cells, with a greater than fourfold reduction in the number of donor cells recovered relative to controls. However, of the cells recovered, those from both *Bcl11b^+/-^* and *Bcl11b^+/+^* donors alike were still almost exclusively of naïve phenotype (**Figure 4b**), although the recovered Bcl11b^+/-^ cells showed their typical modestly elevated CD44 expression (**Figure 4 – figure supplement 1a**). This result was a striking contrast to the results when prethymic precursors, rather than mature naïve T cells, had been transferred in our chimera experiments (**Figure 4a**).

To circumvent the possibility that Bcl11b expression levels could have affected survival differentially during the isolation and transfer procedures, we also performed a similar experiment in which we first transferred mature naïve CD8 cells to wildtype hosts and then deleted one allele of *Bcl11b*. We transferred *Bcl11b^fl/+^Rosa26^CreERT2/+^* or *Bcl11b^+/+^Rosa26^CreERT2/+^* control naïve CD8 T cells into congenically marked hosts and then treated these hosts with tamoxifen beginning the day following transfer. Two weeks following transfer, similar numbers of donor cells of both genotypes were now found in the spleen. The cells newly deprived of one copy of *Bcl11b* were marked by expression of a lox-stop-lox reporter. However, again, CD8 T cells from both donor origins were >97% naïve (**Figure 4c**). In this case, when Bcl11b dosage was reduced only after full maturation, they did not increase CD44 expression compared to controls either (**Figure 4 – figure supplement 1b**). Analysis of T cells recovered from pooled lymph nodes of the same mice also found no significant differences between genotypes (**Figure 4 – figure supplement 1c**). Thus, although reduced Bcl11b expression both expands development of the T_VM_ population and generates an activation-sensitized naïve population, acute Bcl11b reduction is not sufficient to convert mature, naïve CD8 T cells to T_VM_ cells in peripheral homeostasis.

### Bcl11b haploinsufficient cells preferentially generate T_VM_ cells on first emergence from the thymus after birth

To assess when Bcl11b dose reduction does impact the development of virtual memory T cells, we analysed the phenotype of neonatal splenocytes at day 7 of life, just five days after the first T cells are detectable in the spleen, at around day 2 (Le Campion et al., 2002)(**Figure 4d**). Consistent with other reports (Akue et al., 2012; Smith et al., 2018), the CD8 T_CM/VM_ compartment was expanded in neonates relative to adults, forming approximately 30% of the splenic CD8 T cell pool even in wildtype. However, in *Bcl11b^+/-^* neonates, the T_CM/VM_ compartment already formed approximately 60% of the splenic CD8 T cells. This very early preferential expansion of the T_VM_ phenotype pool in *Bcl11b^+/-^* mice suggested that full Bcl11b dose restrains the normal developmental ontogeny of virtual memory T cells.

### Bcl11b haploinsufficient cells are biased toward T_VM_ upon first generation in chimeras

If the naïve peripheral CD8 population in the Bcl11b-haploinsufficient mice was not the source of the expanded T_VM_ cell pool, then the T_VM_ cells could be emerging as such directly from the thymus. We used the *Bcl11b^fl/+^ Cd4-cre^Tg^* conditional heterozygote mice and *Bcl11b^+/+^ Cd4-cre^Tg^* controls as donors in chimeras to test this by following the leading edge of T cell generation in the early weeks after chimeric engraftment. As before, the recipients were busulfan-conditioned to allow full thymic repopulation by donor cells in a T cell replete peripheral environment when they emerge from the thymus. In these recipients, the thymus was efficiently repopulated at week 3 but few if any donor-derived T cells were detectable in the periphery. By week 4, however, donor T cells were evident in the spleen (**Figure 4e**) and their frequencies substantially increased by week 5 (**Figure 4 – figure supplement 1d, e)**. As shown in **Figure 4e**, when donor cells first emerged 4 weeks post engraftment, both donor genotypes made more naïve-phenotype cells than memory phenotype cells. However, at this early timepoint, donor type cells from conditional heterozygote donors were already generating ∼6x as high a fraction of T_VM_ CD8 cells as donors from *Cd4-cre^Tg^* controls. Other aspects of the Bcl11b dose-reduced CD8 T cell phenotype were evident in these first exported cells, including increased CD44 expression and an increased frequency of Ly6C+ cells within the naïve compartment relative to controls (**Figure 4 – figure supplement 1f, g**). The degree of CD44 deregulation and Ly6C expression within the naïve compartment was stable a week later (**Figure 4 – figure supplement 1h, i**), suggesting little if any contribution of peripheral maturation to the phenotype. Thus, the bias toward producing T_VM_ and Ly6C^+^ naïve populations was established in the Bcl11b haploinsufficient cells within a week of their export from the thymus.

### Thymic Bcl11b protein levels are consistent with haploinsufficiency phenotype manifesting at CD8SP stage

To explore how Bcl11b might alter CD8 thymocyte development, we first interrogated how Bcl11b protein levels normally change through intrathymic differentiation stages, using intracellular staining and flow cytometry. Bcl11b protein levels dipped in wildtype mice following both TCR selection stages; i.e. in the post β-selection immature single-positive to CD69^-^ double positive (DP) transition, and from positively selected DP cells to the mature single positive stages (**Figure 5a**). Of note, we used the developmental decrease in CD69 expression to distinguish between less mature (M1) and more mature (M2) CD4 and CD8SP cells, but not CD73 expression, which has been used by others to try to exclude recirculating cells. Its normal regulated expression in intrathymic γδ cells (Coffey et al., 2014; In et al., 2017) and our chimera reconstitution timecourse data indicated that CD73 is not always a faithful indicator of peripheral origin (**Figure 4 – figure supplement 1j**).

**Figure 5.**
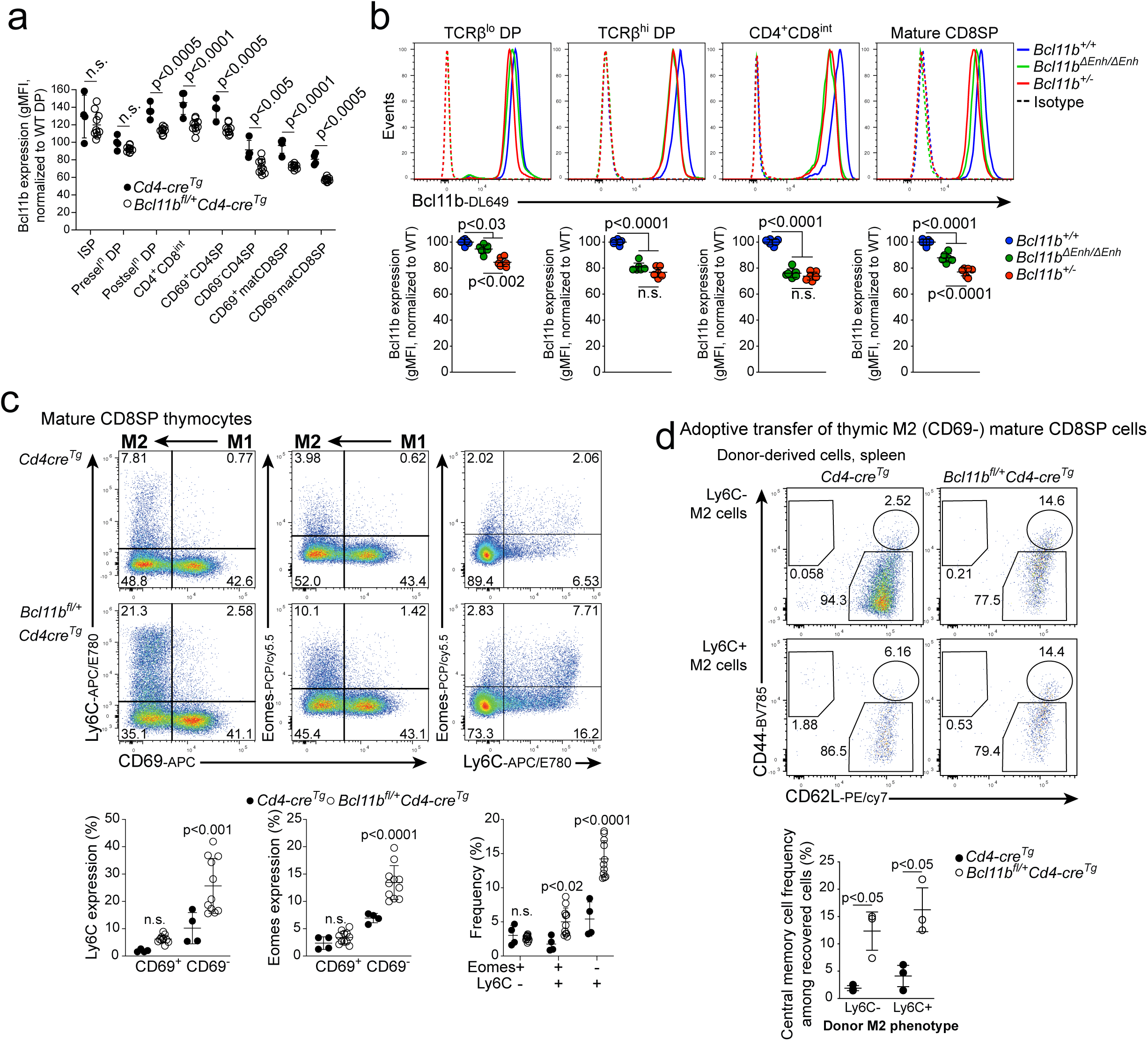
Bcl11b dose in the most mature CD8SP thymocytes determines T_VM_ fate. **a.** Bcl11b protein expression determined by flow cytometry in indicated populations, among conditionally Bcl11b heterozygous or Cre-expressing control thymuses. **b.** Bcl11b protein expression in late thymic developmental stages, comparing co-stained Bcl11b haploinsufficient, enhancer mutant and wildtype control splenic CD8 T cells. Upper, representative histograms, lower, quantification. **c.** Bcl11b conditionally haploinsufficient and Cre-expressing control CD8SP thymocytes were analysed for expression of Ly6C and Eomes among Mature 1 (‘M1’, CD69+) and M2 stages. Upper, representative flow cytometry plots, lower, quantification of marker (co-)expression. **d.** Ly6C+ and Ly6C- M2 stage CD8SP thymocytes from Bcl11b conditionally haploinsufficient or control thymuses were isolated by flow cytometry and adoptively transferred into lymphoreplete hosts. Donor cell phenotype in the spleen analyzed one week later. Upper, representative flow cytometry plots of donor-derived cells’ T_N_, T_CM_ and T_EM_ frequencies, quantified below. Significance tested using one-way ANOVA with Šidák’s (a, c, d) or Tukey’s correction (b).

To test when Bcl11b expression level could be affecting cell fate, we examined how our Bcl11b dose-reduction models affected Bcl11b levels across these thymic stages. That our phenotype of interest manifested with CD4-Cre driven deletion implied that the key difference should occur from the DP stage onward. To measure the modest differences in protein levels between mutant models and wildtype cells accurately, we pooled the corresponding samples from different genotypes in staining them for Bcl11b protein, distinguishing the genetic models by their distinct associated fluorescent proteins (as above). At each stage, a clear difference was seen between the dose-reduction model cells and the controls. In the *Cd4-cre^Tg^ Bcl11b^fl/+^* conditional heterozygote system, where both copies of the gene were intact until the DP stage, the level of Bcl11b protein failed to rise to the levels seen in the controls by the post-selection DP (CD69^+^ TCRβ^+^ DP) and CD4^+^CD8^int^ stages, remaining ∼20% lower than controls in CD69^+^ mature 1 (M1) and CD69^-^ mature 2 (M2) CD8SP cells (**Figure 5a**). Thus, Bcl11b dose could program T_VM_ compartment size at some point from the CD69^+^ TCRβ^+^ DP stage onward.

Across this interval, likewise, the germline heterozygous Bcl11b^+/-^ cells showed approximately 80% as much Bcl11b as the WT controls (**Figure 5b**, left). One question was why the slight decrease in the enhancer mutant could cause any developmental impact at all. Surprisingly, in the TCRβ^hi^ DP to CD4^+^CD8^int^ developmental interval, Bcl11b expression levels in the enhancer mutant transiently stayed as low as in the germline heterozygote model, recovering to restore the WT>enhancer mutant>heterozygote Bcl11b expression hierarchy only by the CD8SP stage (**Figure 5b**). Thus, both *Bcl11b* haploinsufficient and enhancer mutant cells lag in the Bcl11b protein upregulation which normally follows positive selection, notably a stage when CD8 and CD4 SP fates diverge, suggesting that Bcl11b dose may be sensed to affect subsequent mature CD8 T cell fate choice immediately after positive selection.

### Bcl11b dose during positive selection specifically regulates thymic differentiation of virtual memory-like CD8 T cells

The CD8SP populations in the thymuses of *Bcl11b^fl/+^Cd4-cre^Tg^* mice clearly differed from those of Cre control animals. Previous adoptive transfer experiments have identified thymic Eomes-expressing CD8SP cells to be precursors to peripheral virtual memory CD8 T cells in other models (Miller et al., 2020). Ly6C- and Eomes-expressing cells were detectable in the mature CD8SP compartment of both, emerging in the CD69^-^ (M2) population, and they were significantly expanded in the *Bcl11b* conditional heterozygotes (**Figure 5c**). Ly6C^-^ Eomes^+^ cells did not appear to be affected, but the Ly6C^hi^ Eomes^+^ subset was expanded nearly fourfold with Bcl11b dosage reduction, resembling a potential intrathymic version of the T_VM_ cells. Eomes-negative Ly6C^+^ cells were the most numerous expanded subpopulation in the thymic M2 CD8 cells of *Bcl11b^fl/+^Cd4-cre^Tg^* mice relative to Cre controls (**Figure 5c**), resembling the peripheral Ly6C^+^ naïve CD8 T cells.

As Bcl11b dosages were reduced in CD8 and CD4 cells alike, we asked whether thymic CD4SP lineages in these mice might show a similar skewing toward TCR agonist-selected fates, particularly T_REG_ cells. However, we observed no difference between *Bcl11b^+/-^* and littermate control thymuses in the production of T_REG_ lineage precursors (Foxp3^-^CD25^+^GITR^+^ or Foxp3^+^CD25^-^) nor of mature Foxp3^+^CD25^+^ thymocytes (**Figure 5 – figure supplement 1a**), consistent with the unchanged numbers of mature splenic T_REG_ cells described above (**Figure 1 – figure supplement 3c**). Therefore, the main impact of Bcl11b dose reduction in thymic SP cells was seen on the CD8 lineage.

### M2 CD8SP thymocytes, independent of Ly6C expression, include immediate precursors to T_VM_ cells

In prior adoptive transfer experiments (Fig. 4), transfer of pre-thymic precursors (bone marrow chimeras) gave rise to T_VM_ cells, while transfer of mature T cells with reduced Bcl11b expression did not. We therefore tested whether Bcl11b dose has set CD8 T cell fate in the M2 CD8SP thymocytes, by isolating them from *Bcl11b^fl/+^Cd4-cre^Tg^* and Cre-control donors for adoptive transfer into new, wild type hosts. We further fractionated these cells into Ly6C^+^ and Ly6C^-^ donor fractions, to test whether Ly6C specifically marked T_VM_-biased precursors. One week following adoptive transfer, we were able to identify donor-origin CD8 T cells in the spleens and lymph nodes of recipient mice. Bcl11b dose-reduced M2 thymocytes gave rise to T_VM_ cells at significantly higher rates than did Cre-expressing controls (**Figure 5d**).

Despite their enrichment within the Bcl11b haploinsufficient population, naïve cells with Ly6C expression at the time of transfer were not significantly better than Ly6C^-^ cells at acquiring a memory phenotype (**Figure 5d**). Indeed, this was not a longitudinally stable characteristic: control and Bcl11b haploinsufficient cells alike gained or changed Ly6C expression within one week (**Figure 5 – figure supplement 1b**). Thus, the Bcl11b-haploinsufficient M2 CD8SP thymocytes are the immediate precursors to peripheral T_VM_ cells, in a manner independent of Ly6C expression.

### Full Bcl11b dose restricts intrathymic acquisition of a T_VM_ transcriptional profile

We asked how the enhanced bias of Bcl11b haploinsufficient cells to make T_VM_ progeny might already be detected in transcriptomes of cells preparing to leave the thymus. We used flow cytometry to isolate viable M2 mature CD8SP (CD69^-^ TCRβ^+^ CD8^+^ CD4^-^) thymocytes from *Bcl11b^+/-^* and littermate control mice and compared them first by bulk RNA sequencing. Despite the modest changes in Bcl11b protein, 111 genes were upregulated in Bcl11b dose-reduced cells, and 98 genes were wildtype-enriched (adjusted p value < 0.05, 1.5 fold change cutoff, minimum mean expression of 5 FPKM) (**Fig 6a, Supplementary table 4**). GSEA analysis found *Bcl11b^+/-^* M2 thymocytes enriched for signatures of T cell memory (Kaech et al., 2002) and of Ly6C+ naïve CD8 T cells (Jergović et al., 2021) as compared to wildtype M2 cells. As with splenic T cells (**Figure 3g**), a signature of Bcl11b-deficient NK-like cells was also enriched among Bcl11b haploinsufficient cells, including conserved Bcl11b repression targets *Kit, Cd160* and *Cd7* (**Figure 5b**). Thus, a Bcl11b haploinsufficiency phenotype manifests transcriptionally at least by the thymic CD8SP M2 stage.

**Figure 6.**
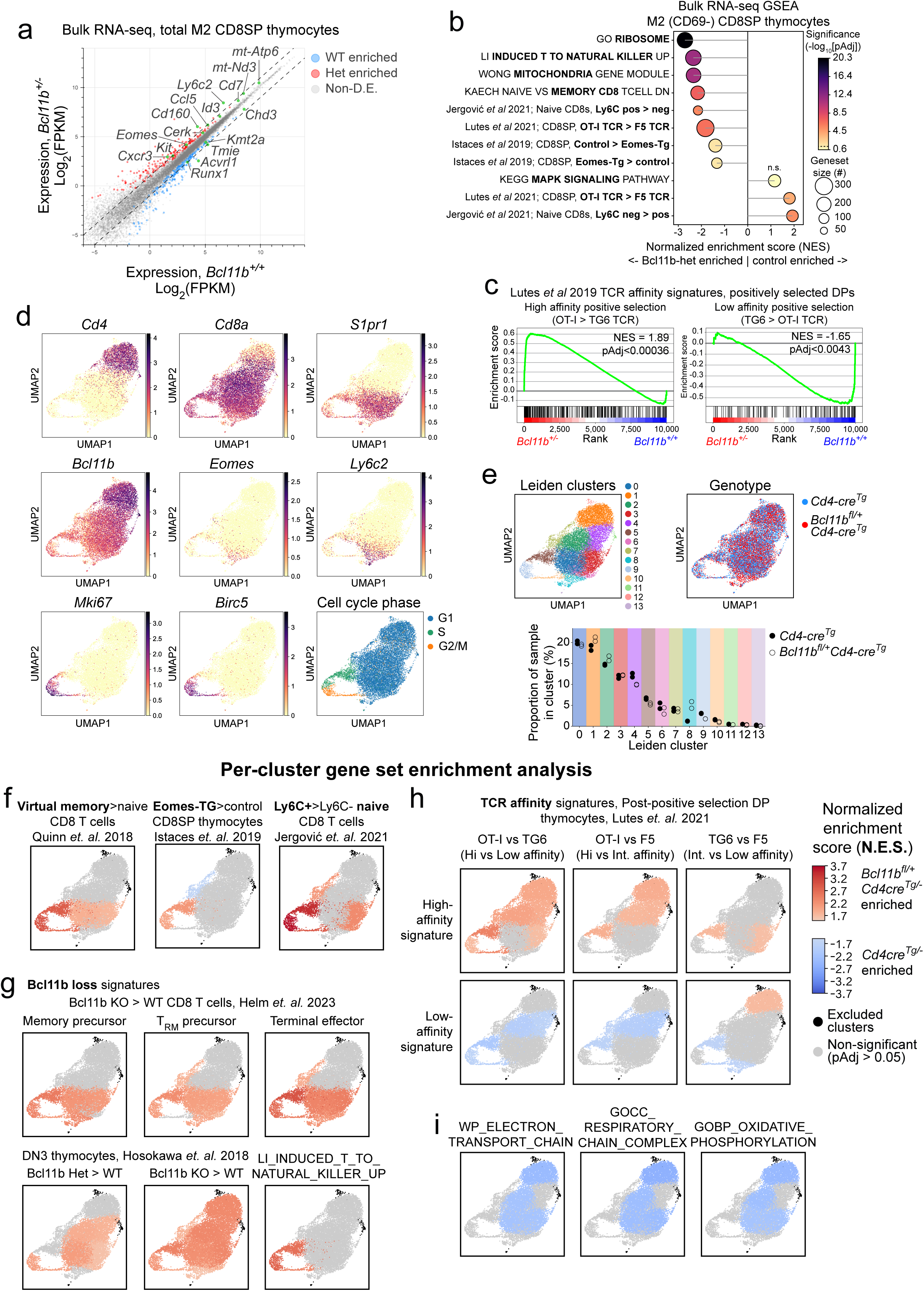
Bcl11b dose activates distinct gene sets found in scRNA-seq and regulates mitochondrial homeostasis in mature thymocytes. **a. - c.** Bulk RNA-seq of Mature 2 stage (‘M2’, CD69-), mature CD8SP (TCRβ+ CD4-CD8+) *Bcl11b^+/-^* and wildtype control thymocytes. **a.** Biplot of mean gene expression in each genotype. Significant genes colored by genotype bias, with select labelled significant genes highlighted in green. Dashed lines indicate a 2-fold change. **b. - c.** Geneset enrichment analysis for indicated gene sets. **d. - i.** Single cell RNA-seq of mature TCRβ+ thymocytes from the CD4^+^CD8^int^ to CD8SP stage comparing *Bcl11b^fl/+^Cd4-cre^Tg^* and Cre-expressing control cells. **d.** Feature plots of expression of indicated genes, and cell cycle score, across a UMAP projection of all included cells. **e.** Upper, distribution of genotypes (left) and Leiden clusters (right) across the UMAP projected cells. Lower, frequencies of each sample’s cells within each cluster. **f. - i.** Pseudobulk differential expression calls using gene set enrichment analysis to compare the two genotypes within each cluster. Whole clusters are colored by normalized enrichment score (N.E.S.) for the indicated gene sets, with warm colors for clusters in which the geneset is enriched in the Bcl11b haploinsufficient cells and cold colors for clusters in which the gene set is enriched in the *Bcl11b^+/+^* control cells. Non-significant clusters are colored grey and clusters insufficiently populated for testing are colored black. **f**. T_VM_, enforced Eomes expression, and Ly6C+ naïve CD8 T cell relevant gene sets. **g.** Bcl11b loss gene sets. **h.** Genesets of DP thymocytes positively selected with transgenic TCRs with a known hierarchy (OT-I>F5>TG6) of self-reactivity. **i.** Mitochondrial oxidative phosphorylation gene sets.

*Eomes* itself was among those genes derepressed in Bcl11b haploinsufficient cells (**Fig 6a, Supplementary table 4**). However, we observed no convincing enrichment of a signature induced by enforced Eomes expression (Istaces et al., 2019) (**Figure 6b**). This suggests that later, Eomes-driven aspects of the memory program may not manifest detectably until after export. We also observed no enrichment for genesets simply indicating the presence vs absence of T cell receptor signalling, e.g. the MAP kinase signalling pathway (**Figure 6b**; cf. **Figure 3g**). Notably, however, genes induced differentially during positive selection by high vs. low affinity TCR interactions were differentially enriched. Bcl11b haploinsufficient CD8SP cells were enriched for signatures of CD8SP thymocytes selected by high affinity OT-I TCR > by low affinity TG6 TCR (Lutes et al., 2021), while controls were enriched for the reverse signature (**Figure 6b**). Furthermore, the Bcl11b heterozygotes expressed genes upregulated early in response to high-affinity driven positive selection, a signature of the genes that differ between positively selected DP cells in OT-I vs TG6 TCR transgenic mice (Lutes et al., 2021). Again, Bcl11b haploinsufficient cells were enriched for the high affinity OT-I signature and controls for the low affinity TG6 signature, particularly clearly for the leading edge genes in each case (**Fig 6c**). Thus, Bcl11b dose may regulate a key subset of the TCR affinity-dependent transcriptional response.

### Locating early transcriptome effects of Bcl11b haploinsufficiency in CD8SP thymocytes at the level of single cells

To determine the order of molecular events through which Bcl11b dosage-reduced CD8SP thymocytes become biased toward the T_VM_ pathway, we carried out single cell RNA-sequencing (scRNA-seq). To cover the whole range of stages over which this pathway could branch off from the wildtype pathway, we isolated mature TCRβ+ CD8^+^ cells ranging from the CD4^+^CD8^int^ to mature CD8SP stages, from *Bcl11b^fl/+^Cd4-cre^Tg^* and Cre control thymuses.

The recovered events ranged from the most immature cells in the continuum (higher *Cd4, Zbtb7b, Ccr9, Cd28, Tcf7, Ets1* and *Tox* expression) to the most mature peri-export thymocytes [more *Cd8a*, *Cd8b1*, *Runx3*, *B2m*, *H2-Q7* (Qa2), *S1pr1* and *Sell* expression]. Cells enriched for expression of *Eomes*, *Ly6c2* (Ly6C), *Il2rb*, *Cxcr3* and *Ccl5*, among other T_VM_-associated genes constituted a small subset among the most mature cells in the pooled sample. A distinct ‘loop’ of cycling (*Mki67*^+^, *Birc5*^+^) cells was also evident among the most mature cells, with the G2/M end terminating adjacent to the T_VM_-like cluster in UMAP space (**Figure 6d**). Leiden graph-clustering showed no single cluster that was unique to either genotype; however, Bcl11b haploinsufficient cells included a threefold higher proportion of T_VM_-like cluster 8 cells than wildtype cells did (**Figure 6e**). Thus, concordant with the ATAC-seq results, Bcl11b heterozygotes traversed essentially the same states as wildtype cells; however, they yielded more T_VM_-like products.

To identify possible drivers or predictors of the differential outputs, we examined when any differences in gene expression emerged between genotypes. We used pseudobulk differential expression analysis to compare the wildtype and Bcl11b haploinsufficient cells within clusters, and also within pooled M1-like clusters (2 and 4), M2-like clusters (0 and 3), and cycling clusters (5, 9 and 10) to increase statistical power. Cluster 1 contained the earliest CD4^+^CD8^int^ cells and cluster 8 the terminal, Eomes^+^ T_VM_-like cells. Clusters preceding cluster 8 showed genotype-linked significant differential expression (Fold change > 1.5, adjusted p value < 0.05) of a few transcriptional/epigenetic regulators (*Lmo4*, *Chd3, Id3*) and other Bcl11b-regulated genes (*March5*, *Izumo1r*, *Auts2*). However, a T_VM_-like gene signature as such (*Ly6c2*, *Ccl5*, *Sidt1*, *Plac8*, *Cd160*) was distinctly differentially expressed between genotypes only within the cycling clusters and cluster 8 (**Supplementary table 6**; violin plots of expression of key differentially expressed and control genes in **Figure 6 – figure supplement 1a**).

As Bcl11b dose appears to regulate the probability that cells will enter the T_VM_ (and/or Ly6C+ naïve) lineage, we asked whether reduced Bcl11b only affected a subset of the population, or whether all the heterozygous cells within a cluster were affected. We identified sets of strongly genotype-associated (Fold change > 2, adjusted p value < 0.05) genes in any pseudobulk comparison (**Supplementary table 6**), and plotted histograms of the WT-enriched and Bcl11b-Het enriched geneset scores for cells in each of our major cluster groups. In each cluster group, the score distributions displayed a broad lateral shift between wildtype and heterozygote samples, but the Bcl11b haploinsufficient pools showed no sign of a mixture of distinct wildtype-like and abnormal cells (**Figure 6 – figure supplement 1b**). Thus, Bcl11b dosage reduction appears to affect transcription across the whole population.

### Bcl11b dosage reduction effects related to other T_VM_ promoting pathways

To compare possible drivers of T_VM_ development in the Bcl11b haploinsufficiency model with those from other models, we tested whether specific pathway gene sets from different hypothesis-relevant models became enriched in Bcl11b-reduced cells, using rank-based GSEA of pseudobulk differentially expressed genes in individual clusters (**Figure 6f-i**; **Figure 6 – figure supplement 2a-b**). As expected, T_VM_ vs Naïve (Quinn et al., 2018), Eomes transgenic vs control CD8SP (Istaces et al., 2019), and Ly6C+ vs Ly6C- naïve CD8 T cell (Jergović et al., 2021) signatures were enriched among Bcl11b haploinsufficient cells, especially in cluster 8 (**Figure 6f**). Notably, the cell cycle clusters not only showed the highest normalized enrichment scores for these gene sets, but were also the only clusters before cluster 8 consistently enriched for all these signatures, suggesting that Bcl11b-reduced cells become biased to a T_VM_ fate during cycling. Various indices of *Bcl11b* reduction in other T-lineage contexts (Helm et al., 2023; Hosokawa et al., 2018; P. Li et al., 2010) were also enriched in Bcl11b haploinsufficient M2 cells (**Figure 6g**), most consistently among the cycling cells. In stark contrast, however, geneset signatures for the cytokine signals elsewhere reported to contribute to T_VM_ or Ly6C+ naïve CD8 T cell differentiation, such as IL-4, IL-15 and type I interferon signalling, showed no consistent enrichment whatsoever in Bcl11b heterozygotes (**Figure 6 – figure supplement 2a**).

These dynamics were tracked (**Figure 6 – figure supplement 2c** (right plots)) by plotting gene set scores against UMAP2 coordinate values, which largely reflect developmental time (top to bottom). Bulk M2 Bcl11b heterozygote-enriched genes began to show genotype-biased expression genotypes in mid-pseudotime, then split from controls in enrichment of T_VM_-associated and finally in Eomes-associated gene set scores in late M2 (UMAP2<-2.5) (**Figure 6 – figure supplement 2c**, middle). In contrast, IL-4 response-associated genesets never became enriched (**Figure 6 – figure supplement 2c**, center).

If Bcl11b dose really controls T_VM_ lineage commitment via influencing the cell-intrinsic interpretation of TCR affinity, differential TCR affinity-associated responses should precede T_VM_ emergence. Again we examined signatures that distinguish DP thymocytes positively selected by the high self-reactivity OT-I TCR from those selected by low (TG6) and intermediate (F5) affinity TCR (Lutes et al., 2021)(cf. **Fig. 6c**). Results confirmed that the high affinity signatures were enriched in Bcl11b haploinsufficient cells (**Figure 6h**, upper), while the lower affinity signatures were enriched in Bcl11b-replete controls (**Figure 6h**, lower). Notably, this enrichment of differential TCR affinity-related gene signatures became detectable in Bcl11b haploinsufficient cells from the earliest population observed (cluster1, CD4^+^ CD8^int^), preceding the enrichment of T_VM_, Ly6C+ naïve, and Bcl11b loss signatures, as well as through the cell cycle clusters. Of note, GSEA analysis identified different groups of genes in the leading edges of differential TCR affinity-associated genes that were enriched in cluster 1 Bcl11b heterozygotes as opposed to the leading edge genes enriched in cluster 5 heterozygotes. The leading edge genes for cluster1 distinguished the genotypes mostly early in the developmental continuum (UMAP2 >5.0, though also in cycling cells). Differential TCR-affinity associated leading edge genes that were enriched in the cluster 5 (early cycling M2 cells) comparison mainly distinguished genotypes later (**Figure 6 – figure supplement 2c**, bottom; UMAP2<2.5); accordingly, these genesets only partially overlapped (**Figure 6 – figure supplement 2d**). Thus, Bcl11b dose mimics the developmental interpretation of differential TCR affinity signals through shifting genes, from the CD4^+^CD8^int^ to the cycling CD8SP thymocyte stage, when T_VM_ program genes are finally activated.

### Chromatin accessibility is minimally affected by Bcl11b haploinsufficiency

To assess how Bcl11b dose reduction exerted these effects, we asked whether Bcl11b haploinsufficiency was altering chromatin accessibility in thymocytes to poise them for future T_VM_ cell fates. We performed assay for transposase accessible chromatin by sequencing (ATAC-seq) on flow cytometrically sorted *Bcl11b^fl/+^Cd4-cre^Tg^* or *Cd4-cre^Tg^* control cells. From two donors (male and female) of each genotype, we processed five populations: M1 CD8SP thymocytes, Ly6C- M2 CD8SP thymocytes, Ly6C+ M2 CD8SP thymocytes, peripheral naïve CD8 T cells and peripheral central memory CD8 T cells. Libraries were highly consistent both between genotypes and between subset replicates (**Figure 7**, **Figure 7 – figure supplement 1a**), with key T lineage loci such as the *Cd3* locus showing a consistent accessibility profile across stages (**Figure 7a**, left), and developmentally-relevant loci such as *S1pr1* noticeably changing from the thymus to the spleen (**Figure 7a**, right).

**Figure 7.**
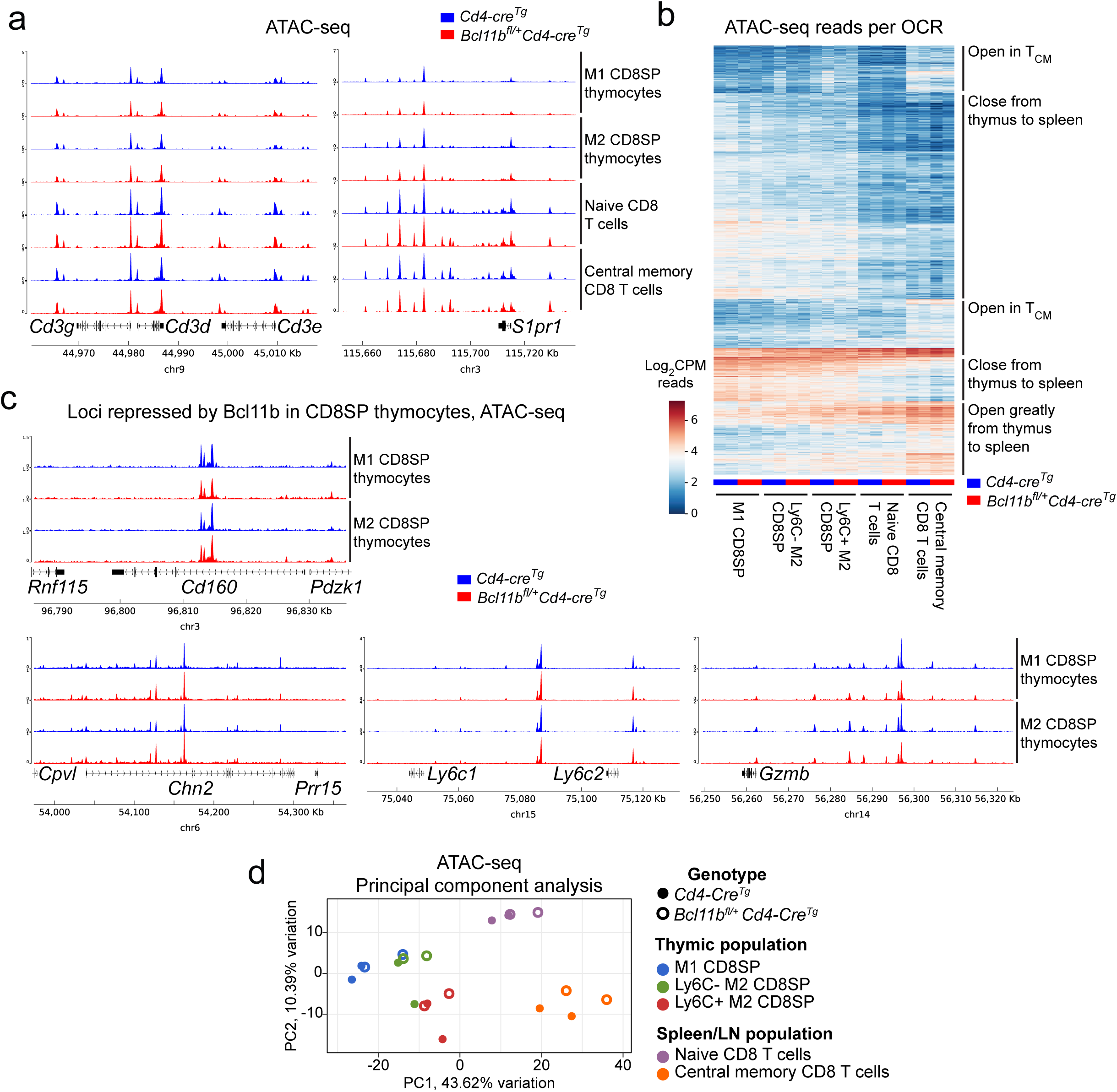
Bcl11b haploinsufficiency alters gene expression without altering chromatin accessibility. **a. - d.** ATAC-seq was performed on thymic M1, Ly6C- M2 and Ly6C+ M2 thymocytes and naïve and central memory splenocytes from *Bcl11b^fl/+^Cd4-cre^Tg^* and Cre-expressing controls. **a.** Replicate-merged genome browser tracks from key loci. **b.** Normalised read counts within called open chromatin regions found to significantly differ between populations. **c.** As in **a**. **d.** Principal component plot of all 20 samples.

Of 71,215 open chromatin regions (OCRs) called across all samples (**Supplementary table 5**), 14,933 were found to be differentially enriched (pAdj < 0.05, fold change > 2) in any pairwise population-population comparison (**Figure 7b**, **Supplementary table 5**). There were clear differences in chromatin opening/closing patterns across developmental stages. However, no difference in accessibility was evident between genotypes (**Figure 7b**), even at loci that are highly upregulated in *Bcl11b* heterozygotes (e.g. *Cd160*, *Ly6c2, Chn2, Gzmb*) (**Figure 7c**). While principal component analysis separated populations, there was no consistent divergence between samples by genotype (**Figure 7d**). Indeed, just two OCRs were identified as significantly different (pAdj < 0.05) between genotypes, in the *Igh* and *Cblb* loci (**Figure 7 – figure supplement 1b**), neither of which are significantly differentially expressed in RNA-seq comparisons. Thus, despite clear shifts in chromatin accessibility across late CD8 T cell differentiation, there was no substantial difference between *Bcl11b* haploinsufficient and control populations at a given stage. Bcl11b dosage affected gene expression without altering OCR patterns.

### Bcl11b dose regulates mitochondrial dynamics of developing CD8 T cells

We next investigated the impacts of the most Bcl11b dosage-affected gene sets. Both intrathymic and peripheral CD8 T cells showed effects of *Bcl11b* genotype on mitochondrial-associated genes, but with differing signatures in peripheral CD8 T cells (**Figure 3g**) than in M2 CD8SP thymocytes. In bulk M2 thymocyte transcriptome comparisons, two mitochondrially encoded genes, *mt-Nd3* and *mt-Atp6*, were significantly upregulated in Bcl11b^+/-^ CD8SP thymocytes (**Figure 6b),** but in our single-cell analyses, mitochondrial genesets overall were consistently biased toward control genotype cells (**Figure 6i**). One gene in particular, *March5 (*alternatively, *Marchf5* or MITOL), is a known regulator of mitochondrial dynamics (Liu et al., 2023; Yonashiro et al., 2006) that was consistently downregulated in the Bcl11b haploinsufficient cells (**Figure 6 – figure supplement 1a**).

To assess what impacts these differences could have on the cells, we compared mitochondrial content and membrane potential by flow cytometry in T cells from *Bcl11b^+/-^* and littermate control mice. In wildtype thymocytes, the maturation from DP to CD8SP cells was associated with substantial reductions in mitochondrial membrane potential, measured by MitoTracker DeepRed FM uptake, despite slight increases in mitochondrial content based on the mitochondrial outer membrane protein Tom20. Notably, whereas Bcl11b haploinsufficiency resulted in a slight decrease in mitochondrial membrane potential among thymic CD4SP cells, among mature CD8SP thymocytes the haploinsufficient cells retained a significantly higher degree of mitochondrial polarization than controls, despite unchanged mitochondrial content (**Figure 8a**). In the periphery, Bcl11b haploinsufficient splenic CD8 T cells still maintained increased mitochondrial membrane potential relative to controls in naïve and T_VM/CM_ subsets (**Figure 8 – figure supplement 1a**), albeit now accompanied by a slight loss of mitochondrial mass (Tom20). Thus, Bcl11b dose directly or indirectly regulates mitochondrial activity and dynamics from the single positive thymic stage onward, which could contribute to T_VM_ accumulation (Liao et al., 2023; Liu et al., 2023).

**Figure 8.**
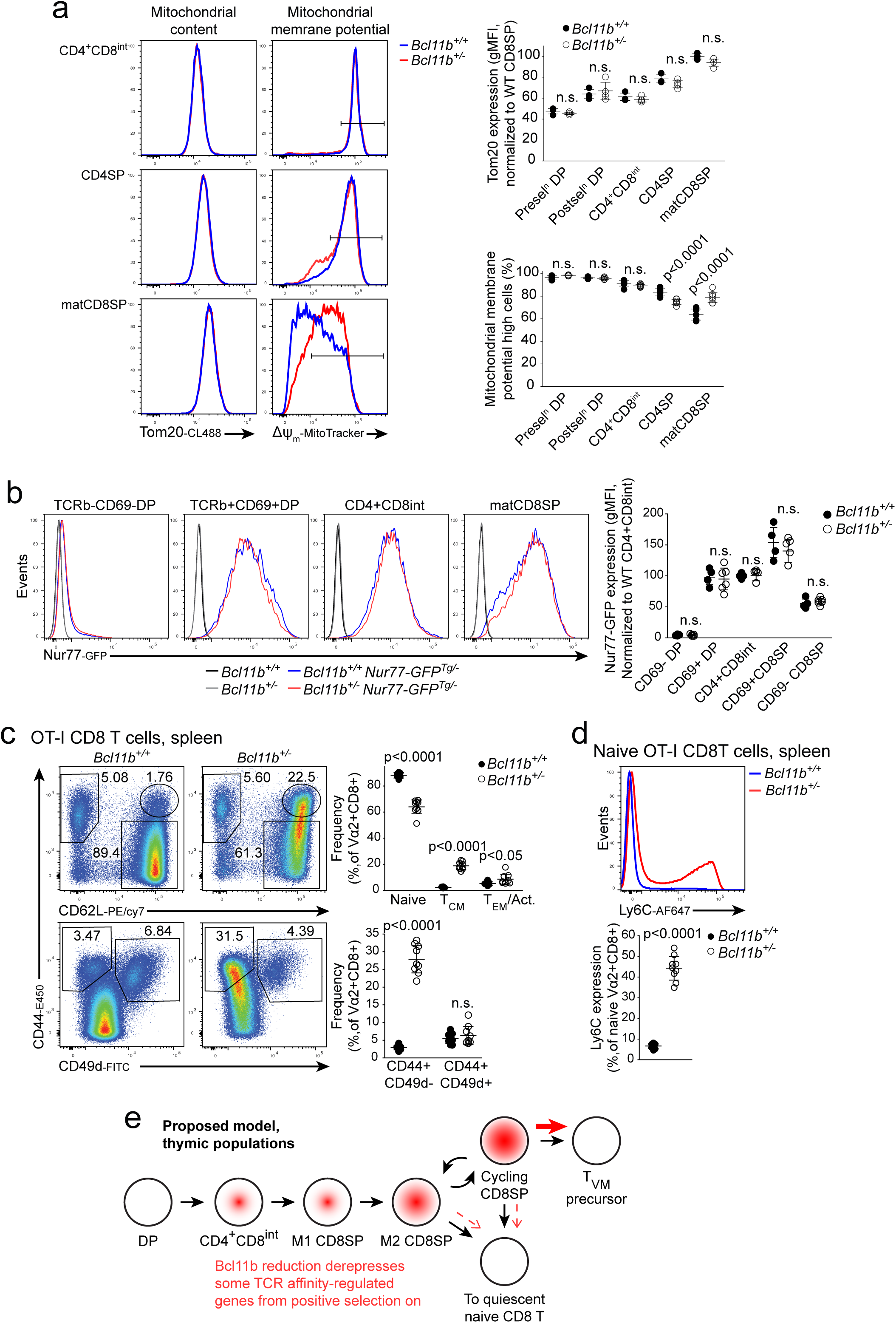
Bcl11b haploinsufficiency phenotype does not result from altered TCR signaling. **a.** Flow cytometric analysis of mitochondrial content and polarization. Representative histograms of Tom20 (left) and MitoTracker DeepRed FM (right) straining in indicated late thymic populations, quantified below. **b.** Representative histograms of Nur77-GFP expression by indicated thymic populations, quantified right . **c. - d.** Flow cytometric analysis of splenic CD8 T cells from OT-I TCR transgenic *Bcl11b^+/-^* and littermate *Bcl11b^+/+^* controls. **c.** Representative flow cytometry plots (left) of, upper, T_N_, T_CM_ and T_EM_ frequencies, and lower, ‘true memory’ vs T_VM_ frequencies, quantified right. **d**. Representative histogram of Ly6C expression by naïve CD8 T cells of each genotype (upper), quantified below. **e.** Model of a proposed intrathymic commitment pathway for T_VM_ cells. Significance tested using one-way ANOVA with Šidák’s correction (a. – c,) or Student’s two-tailed t test (d.).

### Test of Bcl11b dosage effect on history of TCR signalling via a Nur77-GFP reporter

Reduced Bcl11b could enhance TCR-strength-dependent responses through multiple signaling pathways and response systems. If it altered the TCR complex itself, all responses would be affected. If its effects were further downstream, some TCR-signaling responses might not be affected. We crossed the Bcl11b-null allele to mice carrying the *Nr4a1* (Nur77)-GFP reporter, which sensitively reports TCR signal strength in thymocytes and peripheral T cells (Moran et al., 2011), and verified that this reporter showed characteristic differences in GFP expression between pre- and post-selection DP, CD4^+^CD8^int^ and CD8SP thymocytes. However, globally there was little if any difference between the Bcl11b heterozygous and Bcl11b wildtype cells in each compartment (**Figure 8b**). Among M2 CD8SP cells, Nur77 reporter decreased as cells increased in their expression of Ly6C and was lowest in Ly6C^hi^Eomes^+^ cells (**Figure 8 – figure supplement 1b**). However, within these fractions, Nur77 reporter expression was unaffected by Bcl11b haploinsufficiency (**Figure 8 – figure supplement 1c**). Thus, Bcl11b-Het cells in the CD8SP thymocyte compartment showed no evidence that they experienced globally altered TCR signals.

In the periphery, the compartments expanded by Bcl11b haploinsufficiency again expressed Nur77 reporter almost indistinguishably from wildtype controls. Both CD44^high^CD49d^-^ T_VM_ cells (**Figure 8 – figure supplement 1d**) and CD44^low^Ly6C^+^ naïve cells (**Figure 8 – figure supplement 1e**) displayed reproducibly similar Nur77 reporter expression between Bcl11b heterozygous and control spleens. Curiously, Bcl11b haploinsufficiency had some impact on Nur77-GFP expression in the populations not expanded by Bcl11b dose reduction. CD49d^+^ ‘true’ memory cells showed lower Nur77-GFP expression in *Bcl11b^+/-^* spleens than in controls (**Figure 8 – figure supplement 1d**), suggesting they were being maintained on weaker tonic signals than wildtype control memory cells, while haploinsufficient Ly6C^-^ naïve CD8 T cells displayed greater expression of the Nur77 reporter than did *Bcl11b^+/+^* littermate controls (**Figure 8 – figure supplement 1e**), possibly connected with the lower activation threshold in these naïve cells (cf. Fig. 3 – fig. supplement 1e). Taken together, however, these results show that reduced Bcl11b dosage in thymocyte development does not affect TCR signal strength per se but affects particular components of the downstream response to TCR signaling.

### Bcl11b haploinsufficiency drives T_VM_-biased development even when TCR specificity is fixed

*Bcl11b* haploinsufficiency could cause a shift in the affinity spectrum of TCRs effective in positive selection or affect T cell responses by a mechanism independent of TCR affinity itself. To resolve this, we assessed whether enforcing a common TCR affinity would eliminate or exacerbate the Bcl11b haploinsufficiency phenotype. We crossed the *Bcl11b*-null allele into mice transgenic for the class I MHC-restricted OT-I TCR, which mediates relatively strong interaction with self-MHC (Ge et al., 2004; Lutes et al., 2021) but does not encounter its cognate peptide antigen in a normal B6 mouse. Among TCR transgene-expressing Vα2^+^ CD8 T cells in the spleen of these mice, Bcl11b haploinsufficiency still markedly expanded the CD62L^+^CD44^+^ central memory phenotype cells at the expense of naïve T cells when compared with controls (**Figure 8c**, upper), just as in a non-TCR transgenic. Again, the expanded population lacked CD49d expression clearly identifying the cells as T_VM_ (**Figure 8c**, lower). Among naïve OT-I CD8 T cells, Bcl11b haploinsufficiency also resulted in a >5-fold increase in the rate of Ly6C positivity (**Figure 8d**), just as in polyclonal TCR populations. Thus, Bcl11b dose programs mature CD8 T cell fates independent of the strength or diversity of the TCR signals exogenously received.

## Discussion

The transcription factor Bcl11b is indispensable for establishing and maintaining T cell identity, but its expression level varies in different T-cell subsets, and undergoes reversible downregulation during T-cell activation. Here we have shown that <2-fold reduction in Bcl11b expression level at a critical stage in the thymus has a lasting effect on the fates of CD8 T cells that cannot be reproduced by later dosage reduction in peripheral naïve T cells. Bcl11b heterozygosity only modestly impacted thymocyte development overall, but strongly increased a virtual memory T cell-like population in the periphery at the expense of naïve CD8 T cells, from neonatal and adult cohorts of T cells alike, in three different models of dose reduction and one model of a functional hypomorph. The role of Bcl11b that was so dose-sensitive in CD8 T cell development depended at least in part on its corepressor recruitment motif, and the Bcl11b^R3S^ phenotype implied that Bcl11b depends on recruitment of repressive complexes more for its roles in later-stage thymocyte development than for its roles in DN-stage thymocyte development.

The most remarkable feature of this phenotype is how slight a reduction of Bcl11b protein level was sufficient to alter the balance of cell fate choices in CD8 T cell maturation. The key seems to be that Bcl11b levels are reduced during a particular developmental stage that is ultrasensitive to its levels, different from the resulting peripheral T cell populations that show the most obvious alterations. Bcl11b dosage reduction has also been associated with increased production of “innate-like” CD8SP thymocytes in an earlier study (Hirose et al., 2015), where heterozygous deletion of *Bcl11b* exon 2 by CD4-Cre was combined with a missense mutation on the other allele. The resulting disruption was severe enough to reduce positive selection efficiency overall, however, complicating the interpretation. Here, we deleted exon 4, which encodes all the DNA-binding zinc fingers plus a crucial protein-interaction domain, but in the heterozygote Bcl11b levels remained high enough to support general CD4 and CD8 positive selection efficiently. This study, and a recent study of Ets1 (Chandra et al., 2023), indicate how subtle a quantitative change in a critical regulatory factor can generate a selective but durable physiological impact.

Rather than simply presenting an intermediate or ‘partial phenotype’ compared with homozygous Bcl11b loss, here haploinsufficiency manifested distinctly. Broadly, loss of both alleles of Bcl11b is associated with a degree of lineage instability across T cell subtypes (Sidwell and Rothenberg, 2021) with reduced expression of T lineage markers and TCR signalling molecules, and impaired proliferation (Zhang et al., 2010). Here, however, naïve Bcl11b haploinsufficient CD8 T cells showed enhanced sensitivity to TCR stimulation in the periphery and proliferated faster than equivalently stimulated controls. Thus, partial and complete Bcl11b loss appear to result in not just distinct but opposing effects on CD8 T cell biology.

The developmental timing of the Bcl11b haploinsufficiency impact on CD8 T cells described here was established with CD4-Cre dependent deletion and a series of adoptive transfer experiments. M2 CD8SP thymocytes but not peripheral naïve CD8 T cells were identified as the immediate precursors to the expanded peripheral virtual memory T cell compartment. The small difference in Bcl11b level did not result in a clear chromatin accessibility change: in each stage, the accessible chromatin landscapes of Bcl11b haploinsufficient and control cells appeared to be indistinguishable. However, scRNA-seq showed that a small, specific list of genes became differentially expressed by corresponding haploinsufficient and wildtype cells across the CD8SP cell maturation process, especially in the later stages. These specific Bcl11b heterozygote-enriched genes included features of typical T_VM_ precursor differentiation as well as signatures seen in DP cells undergoing positive selection by high-affinity TCR interactions.

What is the relationship of the Bcl11b level-dependent pathway to other pathways that reportedly generate increased numbers of T_VM_ cells? Here we excluded specific hypotheses of how Bcl11b dose regulates T_VM_ differentiation: regulating conversion from naïve cells in the periphery, altering the affinity of TCR interactions, or regulating the transcriptional signature induced by T_VM_-promoting cytokines. There were no changes in expression of regulatory genes whose disruption has been associated elsewhere with T_VM_ differentiation, such as *Dot1l*, *Ezh2*, *Hdac7*, *Dock2* or *Ikzf3*. Thus, the Bcl11b dose-sensitive mechanism is distinct from these. However, we identified robust genotype-linked enrichment for signatures of differential TCR signal strength during positive selection among Bcl11b dose-reduced CD4^+^CD8^int^ and CD8SP thymocytes, indicating that the cells respond as if they had been selected by a stronger TCR interaction. This response did not reflect an increased efficiency of the TCR itself because there was no difference in Nur77 reporter induction between genotypes in defined thymic stages. The phenotype did not require a shift in TCR repertoire, as fixing TCR specificity did not ablate the Bcl11b haploinsufficiency phenotype described here. Instead, our results imply that *Bcl11b* dose reduction increased the interpretation of TCR signal strength through transcriptional de-repression of a subset of TCR-responsive genes, downstream of membrane-proximal events.

Only some TCR affinity-dependent processes were impacted by Bcl11b dose reduction, while others, such as thymic T_REG_ cell differentiation and negative selection, were seemingly unaffected. The ability of the Bcl11b^R3S^ variant, with defective repressor recruitment (Goos et al., 2019), to recapitulate the phenotype suggests that repression was the key role: a sufficient amount of Bcl11b was needed in normal mature CD8SP thymocytes to repress a T_VM_-promoting locus or loci. Taken together, this suggests that Bcl11b dose is controlling T_VM_ pathway access by regulating specific subsets of TCR affinity responsive genes that regulate intrathymic CD8 T cell fate allocation.

Whereas the effect of *Bcl11b* dose reduction on chromatin accessibility was weak, it showed a striking impact on mitochondrial membrane potential during CD8 thymocyte development, sustaining high potential while wildtype cells reduced their mitochondrial potential. This could reflect a generally activation-susceptible state or other causes of high electron transport activity with decreased ATP synthesis (Tovar-Ferrero et al., 2025). A related phenotype was recently reported in human T cells completely deprived of Bcl11b, in which increased fusion actually enhanced mitochondrial function via altered *OPA1* expression (Liao et al., 2023), but *Opa1* was unchanged in murine *Bcl11b* heterozygotes. However, the regulator of mitochondrial division and morphology *March5* (*Marchf5*) was significantly downregulated in M1 and cycling CD8SP cells in *Bcl11b* heterozygotes. As *March5* reduction has been associated with an expanded central memory CD8 T cell compartment in homeostasis (Liu et al., 2023), this is a promising candidate for linking both the T_VM_ and mitochondrial phenotypes of Bcl11b haploinsufficient CD8 T cells. Further work will be needed to explore this.

Finally, there has long been an assumption that late stages of thymocyte differentiation are nonproliferative, although there has been much discussion about the roles of IL-7R and STAT5 signaling in these stages. Our data indicate that at least one cell cycle marks the last intrathymic step of development for many CD8SP cells, and it is here that the greatest transcriptomic impact of Bcl11b dosage reduction was seen. The only cluster-controlled thymic population in which a cohort of T_VM_- and TCR affinity-relevant genes showed consistent enrichment among the Bcl11b haploinsufficient compartment was that of cycling cells. Thus, it is tempting to speculate that proliferation is an inherent component of antigen-independent memory T cell differentiation, and that those CD8SP cells which proliferate prior to export have best access to the T_VM_ fate – possibly linked to division-related dilution of Bcl11b protein itself.

## Author Roles

T.S. conceptualized the project, designed experiments, performed experiments, analyzed data, and wrote the paper. E.V.R. conceptualized the project, designed experiments, analyzed data, wrote the paper, and provided funding.

## Acknowledgments

We gratefully acknowledge Dr. Mark Leid, Oregon State University (present address: Washington State University), for the gift of the Bcl11b^R3S/+^ mice, Dr. Brendan MacNabb for expert guidance in ATAC-seq processing, and former group members Dr. Boyoung Shin (present address: Emory University), for valuable assistance with in vivo cell transfers, and Dr. Hao Yuan Kueh (present address: Yale University), for originally generating the Bcl11b^+/-^ (disruptive mCherry allele) mice. Shirley Pease and Dr. Shoma Nakagawa provided invaluable support in generating and preserving the specialized mouse strains developed here. We also thank Maria Quiloan and Mei Chau for assistance with mouse breeding and genotyping; Ingrid Soto for animal care; Rochelle Diamond, Michael Gregory, Madeline Adolf, Olivia Finney, Diana Perez and Jamie Tijerina for cell sorting; Brian A. Williams of the Single-Cell Profiling and Engineering Center, Caltech, for help with single-cell RNA-seq; and Igor Antoshechkin and Vijaya Rao of the Muriel and Millard Jacobs Genetics and Genomics Laboratory, Caltech, for sequencing; and Diane Trout and Henry Amrhein for sequence analysis system administration. The members of the Rothenberg group are gratefully acknowledged for valuable discussions. This work was supported by a Caltech Baxter postdoctoral fellowship to T. S. and by USPHS grants R01AI135200 and R37AI178114 to E.V.R.

## Data availability

Sequencing data generated in this study have been deposited in the Gene Expression Omnibus, under accession numbers GSE333222, GSE333223 and GSE333224 for bulk ATAC-seq, bulk RNA-seq and single-cell RNA-seq, respectively. The deposited data include raw sequencing files and processed count matrices for all samples analyzed in this study.

## Materials and Methods

### Mice

*C57BL/6J* (JAX 000664) Rosa26-mTomato/mG (JAX 007676), Rosa26-loxp-stop-loxp EYFP (JAX 006148), Cd4-cre (JAX 022071, maintained heterozygously, denoted *Cd4-cre^Tg^* in text) and Cre-ERT2 (JAX 008463) were purchased from the Jackson Laboratory and bred at the California Institute of Technology (Caltech). *Bcl11b^flox^* (Golonzhka et al., 2009) JAX 034469), *Bcl11b^ΔEnh^*, Bcl11b^mCherry^ (Ng et al., 2018), *Bcl11b^mCitrine^* (Kueh et al., 2016) and *Bcl11b^R3S^* (Goos et al., 2019) alleles have been reported previously. *Bcl11b^+/-^* (disruptive mCherry insertion) mice were originally generated at Caltech by H. Y. Kueh and S. Pease by methods used in refs (Kueh et al., 2016; Ng et al., 2018). They were not described fully before but were reported and used for a sample of data shown in (Rothenberg et al., 2016). Both male and female mice were used for this study. All animals were bred and maintained under specific pathogen-free conditions at Caltech according to Institutional Animal Care and Use Committee (IACUC) review and approval.

### Antibodies and Flow Cytometry

Fluorochrome-conjugated antibodies directed against the following mouse antigens were used for analysis by flow cytometry: Bcl11b-FITC (abcam AB123449, clone 25B6, lot 1059198-9), Bcl11b (Bethyl A300-385-A, clone Polyclonal, lot 1 & 2), Anti-Rabbit IgG-DyLight649 (BioLegend 406406, clone Poly4064, lot B325786), B220-Biotin (BioLegend 13-0452-85, clone RA3-6B2, lot 2785846), CD11b-Biotin (BioLegend 101204, clone M1/70, lot B427752), CD11c-Biotin (BioLegend 117304, clone N418, lot B433238), CD122-Biotin (BioLegend 123206, clone TM-β1, lot B316513), CD19-Biotin (BioLegend 115504, clone 6D5, lot B433636), CD3-PCP/Cy5.5 (BioLegend 100218, clone 17A2, lot B365335), CD3-FITC (BioLegend 100204, clone 17A2, lot B415733), CD4-PEcy7 (BioLegend 100528, clone RM4-5, lot B391737), CD4-PE (BioLegend 100512, clone RM4-5, lot B335844), CD4-Biotin (BioLegend 100404, clone GK1.5, lot B381361), CD44-A488 (BioLegend 103016, clone IM7, lot B354292), CD44-Biotin (BioLegend 103004, clone IM7, lot B266682), CD49d-PE (BioLegend 103608, clone R1-2, lot B307588), CD49d-APC (BioLegend 103622, clone R1-2, lot B374373), CD62L-APC (BioLegend 104412, clone MEL-14, lot B355945), CD62L-PE (BioLegend 104408, clone MEL-14, lot B322380), CD62L-A488 (BioLegend 104420, clone MEL-14, lot B370610), CD62L-PEcy7 (BioLegend 104418, clone MEL-14, lot B417092), CD69-APC (BioLegend 104514, clone H1.2F3, lot B361544), CD73-PEcy7 (BioLegend 127224, clone TY/11.8, lot B384644), CD8-APC (BioLegend 100712, clone 53-6.7, lot B365419), CD8-Biotin (BioLegend 100704, clone 53-6.7, lot B427538), GFP-AF488 (BioLegend 338008, clone FM264G, lot B399978), Ki67-A647 (BioLegend 652408, clone 16A6, lot B202257), Kit-PE (BioLegend 105808, clone 2B8, lot B402001), Ly6C-A647 (BioLegend 128010, clone HK1.4, lot B245517), Ly6C-PE (BioLegend 128008, clone HK1.4, lot B239362), NK1.1-APC (BioLegend 108710, clone PK136, lot B229690), Streptavidin-PCP/Cy5.5 (BioLegend 405214, lot B361097), TCRVα2-FITC (BioLegend 127805, clone B20.1, lot B423056), TCRVα2-APC (BioLegend 127810, clone B20.1, lot B433799), TCRβ-Biotin (BioLegend 109204, clone H57-597, lot B381582), TCF1-A488 (Cell Signaling Technology 6444S, clone C63D9, lot 12), CD122-APC (eBioscience 17-1222-82, clone TM-b1, lot E15781-105), CD45-PEcy7 (eBioscience 25-0451-82, clone 30-F11, lot 2972629), CD69-PE (eBioscience 12-0691-81, clone H1.2F3, lot E01331-1631), CD8-PE (eBioscience 12-0081-85, clone 53-6.7, lot E01039-1635), TCRβ-APC (eBioscience 17-5961-82, clone H57-597, lot E07345-1634), TCRβ-APC/E780 (eBioscience 47-5961-82, clone H57-597, lot E08478-1642), CD25-APC/E780 (Invitrogen 47-0251-82, clone PC61.5, lot 2114190), CD4-E450 (Invitrogen 48-0041-82, clone GK1.5, lot 3123623), CD44-E450 (Invitrogen 48-0441-82, clone IM7, lot 2702081), CD49d-FITC (Invitrogen 11-0492-82, clone R1-2, lot 2251589), CD62L-E450 (Invitrogen 48-0621-82, clone MEL-14, lot 2455897), CD8-APC/E780 (Invitrogen 47-0081-82, clone 53-6.7, lot 2944634), Eomes-APC (Invitrogen 17-4875-82, clone Dan11mag, lot 2565474), Foxp3-PE (Invitrogen 12-5773-82, clone FJK-16s, lot 2176028), Foxp3-E450 (Invitrogen 48-5773-82, clone FJK-16s, lot 219559), GR-1-Biotin (Invitrogen 13-5931-86, clone RB6-8C5, lot 2820773), IFNg-APC (Invitrogen 17-7311-82, clone XMG1.2, lot 2410273), KLRG1-APC/E780 (Invitrogen 47-5893-82, clone 2F1, lot 4346707), Ly6C-APC/E780 (Invitrogen 47-5932-82, clone HK1.4, lot 2209835), NK1.1-Biotin (Invitrogen 13-5941-85, clone PK136, lot 3077331), NKG2D-PE (Invitrogen 12-5882-82, clone CX5, lot 2400615), NKG2D-PEcy7 (Invitrogen 25-5882-82, clone CX5, lot 2264902), Rabbit IgG isotype control (Invitrogen 14-4616-82, clone RbNP15, lot 2389680), Streptavidin-A488 (Invitrogen S11223, lot 2480092), Streptavidin-PE (Invitrogen 12-4317-87, lot 2618941), Tbet-PEcy7 (Invitrogen 25-5825-82, clone 4B10, lot 2338686), TCRβ-E450 (Invitrogen 48-5961-82, clone H57-597, lot 2915344), TNFα-E450 (Invitrogen 48-7321-82, clone MP6-XT22, lot 2516616), γδTCR-Biotin (Invitrogen 13-5711-85, clone GL-3, lot 2624391), γδTCR-E450 (Invitrogen 48-5711-82, clone GL-3, lot 2492149), Tom20-CoraLitePlus 488 (Proteintech CL488-80501, lot 210072).

For flow cytometric analysis or isolation, primary tissues were disaggregated to single cell suspensions through a mesh filter (Corning 352350) and incubated for at least 20 minutes in 2.4G2 hybridoma supernatant (produced in house) on ice. Surface antibody staining was performed in CBH buffer (HBSS with 0.5% BSA and 10 mM HEPES) for 30 minutes on ice protected from light. Cell viability was marked using fixable viability dyes (Thermo Fisher L34990 and 65-0866-14, Biolegend 423101) stained in PBS for 15 minutes on ice protected from light, or with propidium iodide (Sigma P4170) or sytox blue (Invitrogen S34857) added directly to samples prior to analysis. Sorted cells were collected into complete RPMI (RPMI media (Thermo-Fisher) supplemented with fetal calf serum (Sigma), non essential amino acids (Gibco), 1mM sodium pyruvate (Gibco), penicillin/streptomycin/glutamate supplement (Gibco) and 55uM β-mercaptoethanol (Sigma)) and kept at refrigeration temperature until further processing. For assessing mitochondrial membrane potential, cells were incubated in MitoTracker Deep Red FM (Thermo Fisher M22426) diluted in complete RPMI at 37°C for 20 minutes.

For intracellular staining of transcription factors, cells were fixed using the eBioscience transcription factor staining buffer set (00-5523-00). For intracellular staining of untethered reporters, cells were pre-fixed for 30 min in 4% paraformaldehyde (Electron Microscopy Sciences) in PBS, prior to any cytokine or transcription factor fixation. For chemokine receptor staining, cells were incubated at 37 °C in the presence of antibody for 1 hour. Flow cytometry data were collected on the MACSQuant (Miltenyi) and CytoFlex-S (Beckman Coulter), and cells flow cytometrically sorted using FACSAria IIu, FACSAria Fusion (BD), CytoFlex-SRT (Beckman Coulter) or SY3200 (Sony) platforms. Flow cytometry data was analyzed using FlowJo software (FlowJo LLC).

### Fine quantitation of Bcl11b expression

To allow direct comparison of Bcl11b content between multiple genotypes under identical conditions, we mixed them for staining in a single tube, taking advantage of the fluorescent reporters intrinsic to each cell type (i.e. the mCherry in the Bcl11b^+/-^ and the IRES-driven mCitrine reporter in *Bcl11b^ΔEnh/ΔEnh^* cells). These were co-stained with reporter-negative wildtype controls, allowing measurement of relative Bcl11b staining intensity among CD8 T cells by flow cytometry by gating on reporter expression.

### Cell purification by magnetic bead depletion

Cells were incubated for at least 20 minutes in 2.4G2 hybridoma supernatant (produced in house) at ≤ 60*10^6^ cells/ml to block Fc receptors, and surface stained with biotin-conjugated antibodies against depletion targets. After washing, cells were then incubated at the same concentration in a 1:10 solution of anti-biotin microbeads (Miltenyi) in HBSS (Gibco) with 2.5mg/ml BSA () and 10mM HEPES (Gibco) for 30 minutes on ice with occasional agitation. Particularly large samples were set on a magnetic rack and the unbound fraction collected three times sequentially to pre-deplete. The cell/bead slurry was passed through an LS column on a Midi MACS magnet (Miltenyi) and the flow-through collected. For naïve CD8 T cells, CD44, CD4, γδTCR, NK1.1, CD19, CD11b, and CD11c antibodies were used. For predepletion of other thymocytes to enrich for thymic CD8SP cells, CD4, γδTCR, NK1.1, CD19, CD11b, and CD11c antibodies were used. A fluorophore conjugated CD4 antibody was present at the same concentration as the biotin conjugated CD4 to allow subsequent identification of CD4-negative cells by flow cytometry, as subsequent staining is blocked.

### Bone marrow and fetal liver chimera generation

Bone marrow chimeric mice were generated by intravenous injection into Busulfan-conditioned hosts, to assay development of the donor cells in an environment replete with mature lymphocytes, according to an established protocol (Montecino-Rodriguez and Dorshkind, 2020). Briefly, adult mice were conditioned with two doses of 20 mg/kg busulfan daily for two days to achieve myeloablation. The following day, donor bone marrow or fetal liver cells were injected retro-orbitally. Animals were left for at least 4 or 8 weeks to allow for reconstitution of thymic or peripheral T cell compartments, respectively.

### Adoptive transfers

Indicated T cell populations were washed and resuspended in PBS, and retro-orbitally injected into congenically distinct host mice. For cells with tamoxifen inducible Cre-ERT2, host mice were injected intraperitoneally with 100mg/kg tamoxifen at 1, 3 and 5 days post transfer. Host mice were analysed for donor-derived cells at the indicated number of weeks post transfer.

### Cell Culture

All cells were cultured in complete RPMI. Naïve (CD62L+CD44-) CD8 T cells were purified using flow cytometric sorting or magnetic bead depletion. Cells were then stimulated by culture in plates pre-coated with anti-CD3 (clone 2C11, 10 μg/mL or as otherwise annotated) and with soluble anti-CD28 (clone 37.51, 2 μg/mL) (Biolegend) and recombinant human IL-2 (1000 IU/ml, Peprotech). For homeostatic conditions, cells were cultured with 20ug/ml IL-2.

Stimulation for cytokine production was performed using phorbol 12-myristate 13-acetate (PMA, 50 mg/mL, Thermo J63916 lot Y221004) and ionomycin (0.5 mg/mL, Invitrogen 124222, lot 2544410) in the presence of Brefeldin A (Sigma B7651, lot 142470) and TNF-α maturation inhibitor 1 (‘TMI-1’, 5 μM, metalloproteinase inhibitor to prevent CD62L loss, Sigma P20336, lot 177094) in complete IMDM for 3 h at 37 °C. The BD Bioscience Cytofix/Cytoperm kit (554714) was used for intracellular analysis of cytokines according to manufacturer’s instructions.

### ATAC-seq

Flow cytometry purified populations were processed for ATAC-seq as described previously (Buenrostro et al., 2015). Briefly, crude nuclear prep was achieved by resuspension in lysis buffer (10 mM Tris-HCl, pH 7.4, 10 mM NaCl, 3 mM MgCl2, 0.1% IGEPAL CA-630) and immediate centrifugation. Nuclei were resuspended in transposition mix (1× TD buffer (Illumina 15027866), 33% PBS, 0.01% digitonin, 0.1% Tween-20, and Tn5 transposase (Illumina 15027916) in UltraPure H2O (Invitrogen 10977-015)) at 1,000 cells/μL and incubated at 37°C for 30 minutes in a thermocycler. Following transposition, DNA clean was immediately performed using a DNA clean and concentrator kit (Zymo D4014) according to the manufacturer’s protocol. For library construction, DNA was again purified using the Zymo DNA clean and concentrator kit following adapter ligating PCR amplification, as described previously (Buenrostro et al., 2015). Small nucleosome-free, 1-, and 2-nucleosome fragments were then enriched using double-sided size selection (0.6X SPRI and 1.8 X SPRI) using AMPure XP beads (Beckman Coulter A63881). DNA concentration was assessed by Qubit HD DNA assay (Invitrogen Q32854), and molarity was calculated by Bioanalyzer HS DNA kit (Agilent NC1738319). After library preparation, sequencing was performed with paired-end sequencing of 2x50 base pair, a targeted depth of 20 million reads per sample on Illumina NextSeq 2000 at the Caltech Genomics Facility. Raw sequencing files and processed count matrices for all samples analyzed in this study were deposited in Gene Expression Omnibus (GEO) under accession number GSE333222.

### Bulk RNA-seq

Total RNA was isolated from 150,000 ∼ 300,000 sorted thymic CD8SP or pooled splenic and brachial, axial and lymph node peripheral CD8 T cell populations using the RNeasy Mini Kit (Qiagen #74105) with on-column DNase I treatment (Qiagen #1023460) according to the manufacturer’s protocol. Sequencing libraries were constructed at the Caltech Genomics Facility and sequenced on Illumina NextSeq (2x50 base pair, targeted to 30 million paired-end reads per sample). Raw sequencing files and processed count matrices for all samples were deposited in GEO under accession number 333223.

### scRNA-seq of thymocytes

Single-cell RNA-seq was performed as described previously (Zhou et al., 2022). TCRβ+ thymocytes in a continuum from the CD4^+^CD8^int^ to CD8SP were isolated by flow cytometry by FACSAria Fusion at the Caltech Flow Cytometry Facility. After isolation, different unique hashtag-oligo (HTO) antibodies (TotalseqA HTO1-HTO11, clones M1/42; 30-F11, BioLegend #155801, #155803, #155805, #155807, #155809, #155811, #155813, #155815, #155817, #155819, #155821) were added to distinguish the identity of each sample and aid in defining doublets. Afterwards, samples were washed with 1×HBSS supplemented with 0.5% BSA and 10 mM HEPES and resuspended to 1x10^6^ cells/1 mL concentration. Then, 20,000 cells were loaded into a 10X Chromium lane, and prepared according to the Chromium v3.1 manual. Single-cell RNA-seq cDNA libraries were prepared using 10X Chromium 3’ capture v3.1 kit. Single-cell hashtag oligo libraries were prepared in accordance with the BioLegend TotalseqA guide. Constructed single-cell RNA-seq cDNA libraries were sequenced to a mean depth of 62,843 reads per cell and hashtag oligo libraries were sequenced for 2,000-2,500 reads per cell on the NextSeq 2000 at the Caltech Genomics Facility. A total of 18,112 cells were captured across four samples (male and female of each genotype), capturing a median of 2,042 genes per cell. Following QC (excluding doublets, high mitochondrial read events and events with insufficient reads or genes) and assigning sample IDs by hashtag oligo labelling, 14,445 singlet events remained, 54% control and 46% Bcl11b haploinsufficient. Raw sequencing files and processed count matrices for all samples were deposited in GEO under accession number 333224.

### Bioinformatic analysis

#### Bulk RNA-seq

The sequencing alignment and gene expression calculation were performed by following the ENCODE RNA-seq pipelines. Briefly, the adapter trimmed reads were aligned to the mouse reference genome GRCm38/mm10 using STAR (v 2.5.1b, (Dobin et al., 2013)), and gene expression was calculated with RSEM (v 1.2.31, (Li and Dewey, 2011)). Then, significant changes in gene expression were determined using DEseq2 (v 1.34.0, (Love et al., 2014)). For the DEseq2 input matrix, low-count genes were filtered out by keeping only those for which the sum of reads all samples was greater than 10.

Differentially expressed genes (DEGs) were defined by average FPKM (across compared samples) > 5, adjusted p-value < 0.05, and absolute Log2 fold-change > 0.585 (at least 1.5-fold increased or decreased).

Genes included for Z-score heatmaps were selected based on indicated differential expression results, with Z-score calculated from FPKM values of individual samples. Heatmaps plotted using pheatmap v 1.0.12.

Gene set enrichment analysis was performed using fgsea (v1.20.0 (Korotkevich et al., 2021)) from DESeq2 output generated as described above. Genes with an average FPKM >5 were used, the Wald statistic used for ranking, and genes without mapped identifiers dropped. Enrichment ranking was performed with default settings, and a minimum pathway size of 50 genes. The molecular signatures database (MSigDB) was queried for gene sets to test against (Castanza et al., 2023; Subramanian et al., 2005). For testing against gene sets generated in-house (including DESeq2 calls of published datasets reanalysed) ENSEMBL identifiers were used.

Principal component analysis was performed using PCAtools v2.6.0 and DESeq2 v 1.34.0

### Single cell RNA-seq

The raw reads from cDNA libraries were aligned to the mouse reference genome GRCm38/mm10 using CellRanger (v 6.1.2) and hashtag oligo libraries were quantified and demultiplexed using in-house tools as described previously (Zhou et al., 2022).

For quality control (QC) and downstream analysis, SCANPY (v1.9.5 (Wolf et al., 2018)) was used. Briefly, for QC, only those cells assigned a unique hashtag oligo identity to exclude doublets, and expressing 800-7000 genes, and with less than 5% of reads mitochondrial were considered, and only those genes with expression detected in at least 10 cells. Inbuilt SCANPY cell cycle scoring was calculated using a cell cycle gene list derived from (Tirosh et al., 2016), while all other geneset scoring utilised the SCANPY scoregenes tool. Pseudobulk differential expression calling was performed using decoupleR v2.1.4 (Badia-I-Mompel et al., 2022) and pydeseq2 v0.5.4 (Muzellec et al., 2023). Gene set enrichment analysis of pseudobulk results was performed using gseapy v1.1.11 (Fang et al., 2023).

### ATAC-seq

Sequenced reads (whether in-house or downloaded public datasets) were mapped to the mouse reference genome GRCm38/mm10 using Bowtie2 (v3.5.1 (Langmead and Salzberg, 2012)). PCR duplicates were removed using Picard (v 2.18.29). Low-quality reads and reads mapped to blacklisted regions and the mitochondrial genome were filtered using Samtools (v 1.9 (Li et al., 2009)). Peak calling was conducted using Genrich (v 0.6) and a unified peak set generated by iteratively merging all peak lists. Briefly, for two summits within 250 bases the less significant peak was dropped until no overlaps remained, and regions of ±250 around each remaining summit constituted the final unified peak list.

Deeptools v3.5.1 (Ramírez et al., 2014) multiBamSummary was used calculate read depths within called regions for aligned libraries. Differential region enrichment (Supplementary table 5) was performed using DESeq2 1.34.0 inputting raw reads per region. Peak regions with an adjusted p value below 0.05 were considered significantly differentially enriched. Principal component analysis was performed using PCAtools v2.6.0 and DESeq2 v 1.34.0, on the 50% most variant peak regions.

### Statistics

Statistical comparisons were calculated with *Prism* (Graphpad) or SciPy (Virtanen et al., 2020) software. Student’s t test (two tailed), one-way ANOVA with Šidák’s or Tukey’s corrections for multiple comparisons, two-way ANOVA with Tukey’s test, or DESeq2 were used as appropriate. For each figure panel, the number of replicates is shown in the figure and the statistical tests used are given in the legends.

---

**Figure 1 – figure supplement 1.**
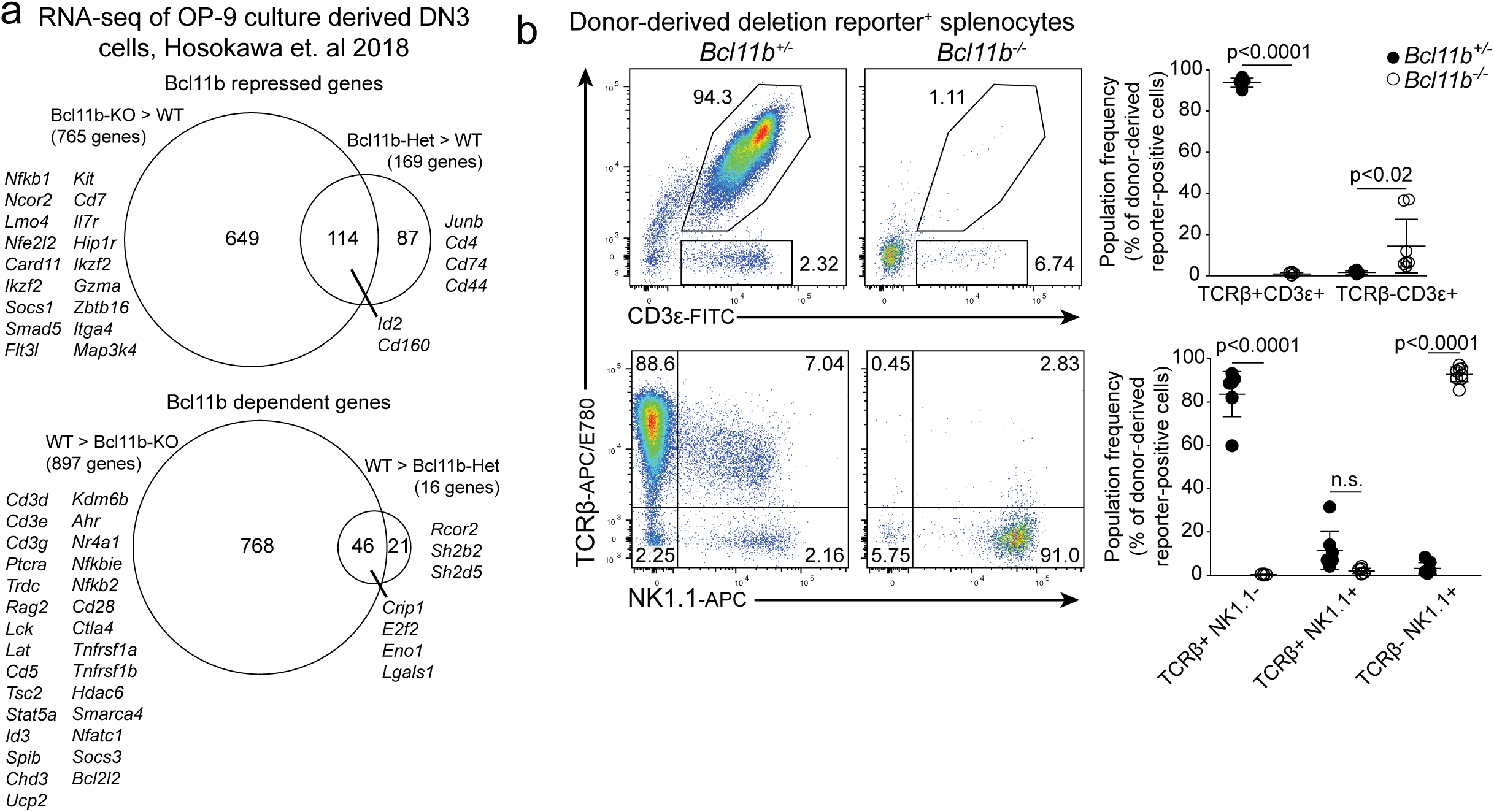
**a.** Reanalysis of differentially expressed genes between homozygous or heterozygous *Bcl11b-*deleted DN3 thymocytes and Cre-expressing controls (Hosokawa 2018). Scaled venn diagrams indicate number of differentially expressed genes, with select genes labelled. **b.** Flow cytometric analysis of fetal liver chimeras reconstituted with homozygous or heterozygous *Bcl11b*-null cells. Right, representative flow cytometry plots of donor-derived cells in the spleen, quantified right. Significance tested using one-way ANOVA with Šidák’s test for multiple comparisons (b).

**Figure 1 – figure supplement 2.**
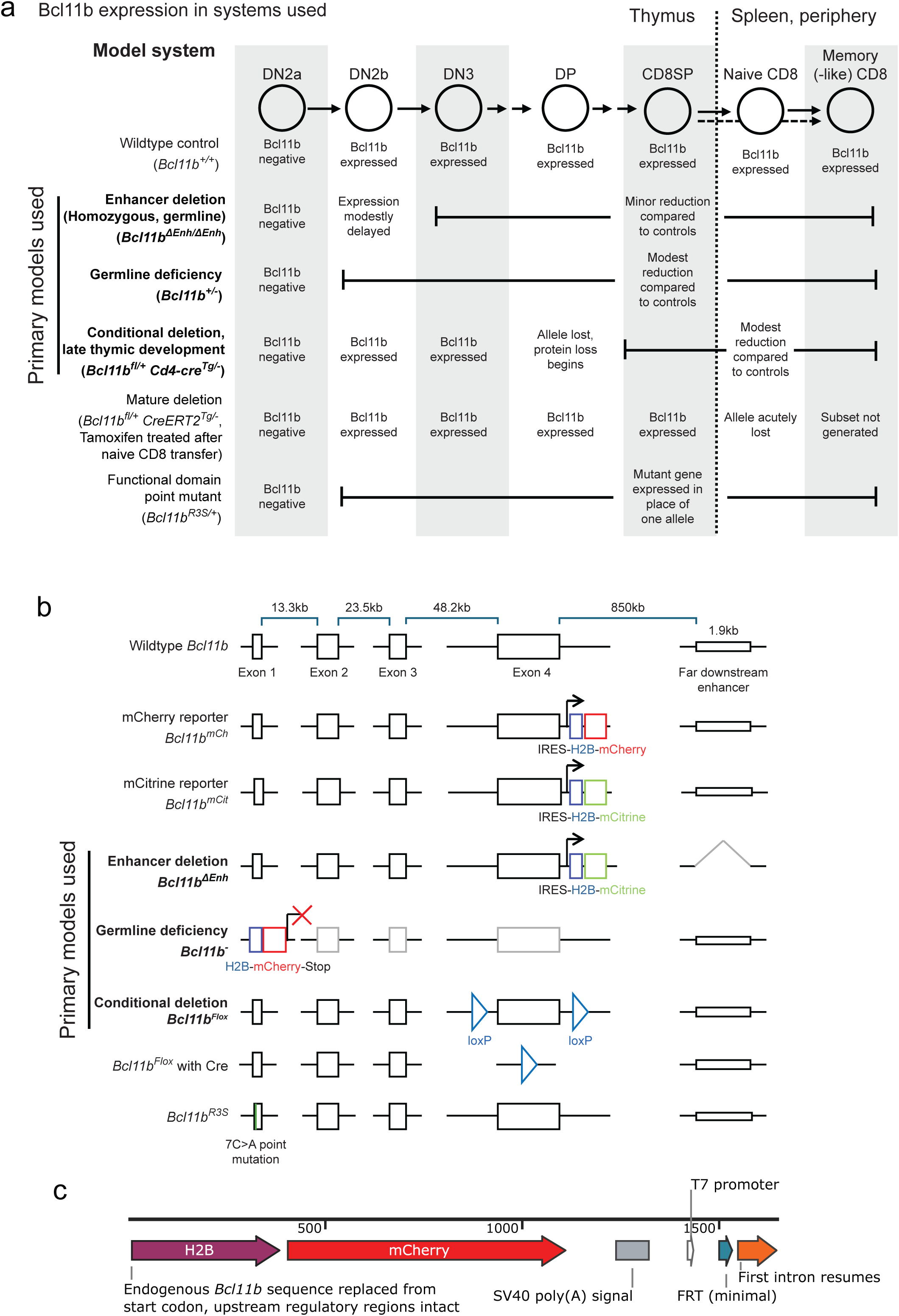
Summary of genetic models used. **a.** The developmental stage in which indicated models display altered Bcl11b expression. **b.** Modifications to the *Bcl11b* locus in each genetic model. **c.** Map of the Bcl11b-null mCherry reporter used here.

**Figure 1 – figure supplement 3.**
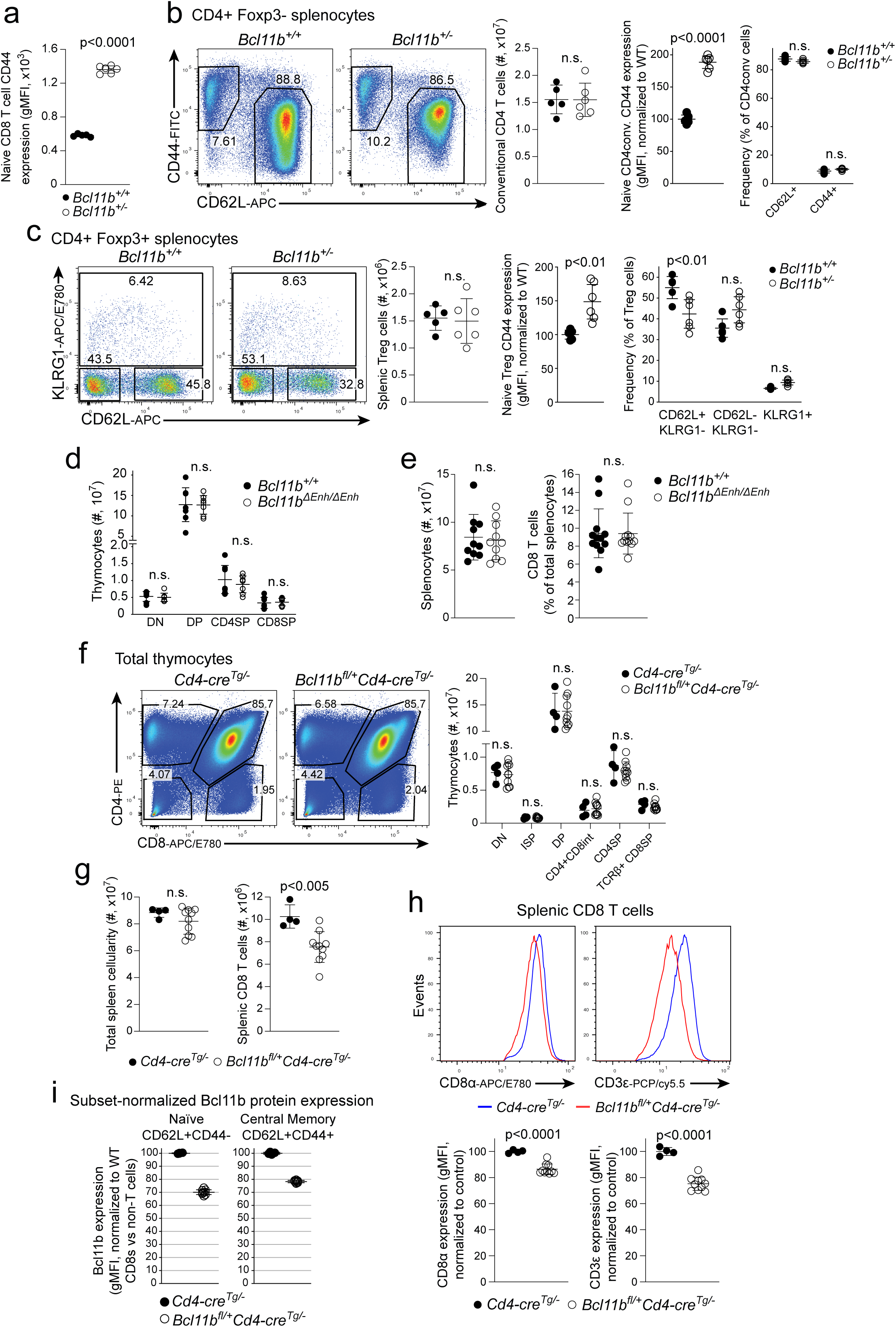
**a.** Quantification of naïve CD8 T cell CD44 expression from Figure 1b. **b.-c.** CD4 T cell compartments of *Bcl11b^+/-^* and littermate control spleens analysed by flow cytometry for activation status and naïve cell CD44 expression. **d.-e.** Flow cytometric characterisation of *Bcl11b^ΔEnh/ΔEnh^* enhancer mutant and control thymic and splenic populations. **d.** Numbers of major CD4/CD8 expressing populations. **e.** Splenic cellularity and CD8 T cell frequencies. **f. - i.** Flow cytometric characterization of *Bcl11b^fl/+^Cd4-cre^Tg^* and Cre-expressing control tissues. **f.** Representative flow plots (left) and quantification (right) of major CD4/CD8 expressing thymic populations. **g.** Splenic cellularity and CD8 T cells numbers of each genotype. **h.** CD8 and CD3 expression level among splenic CD8 T cells of each genotype. **i.** Bcl11b protein content data from Figure 1f normalized to control cells within T_N_ and T_CM_ populations. Significance tested using one-way ANOVA with Šidák’s test for multiple comparisons (b-d, f) or Student’s two-tailed t test (a-c, e, g-i)

**Figure 3 – figure supplement 1.**
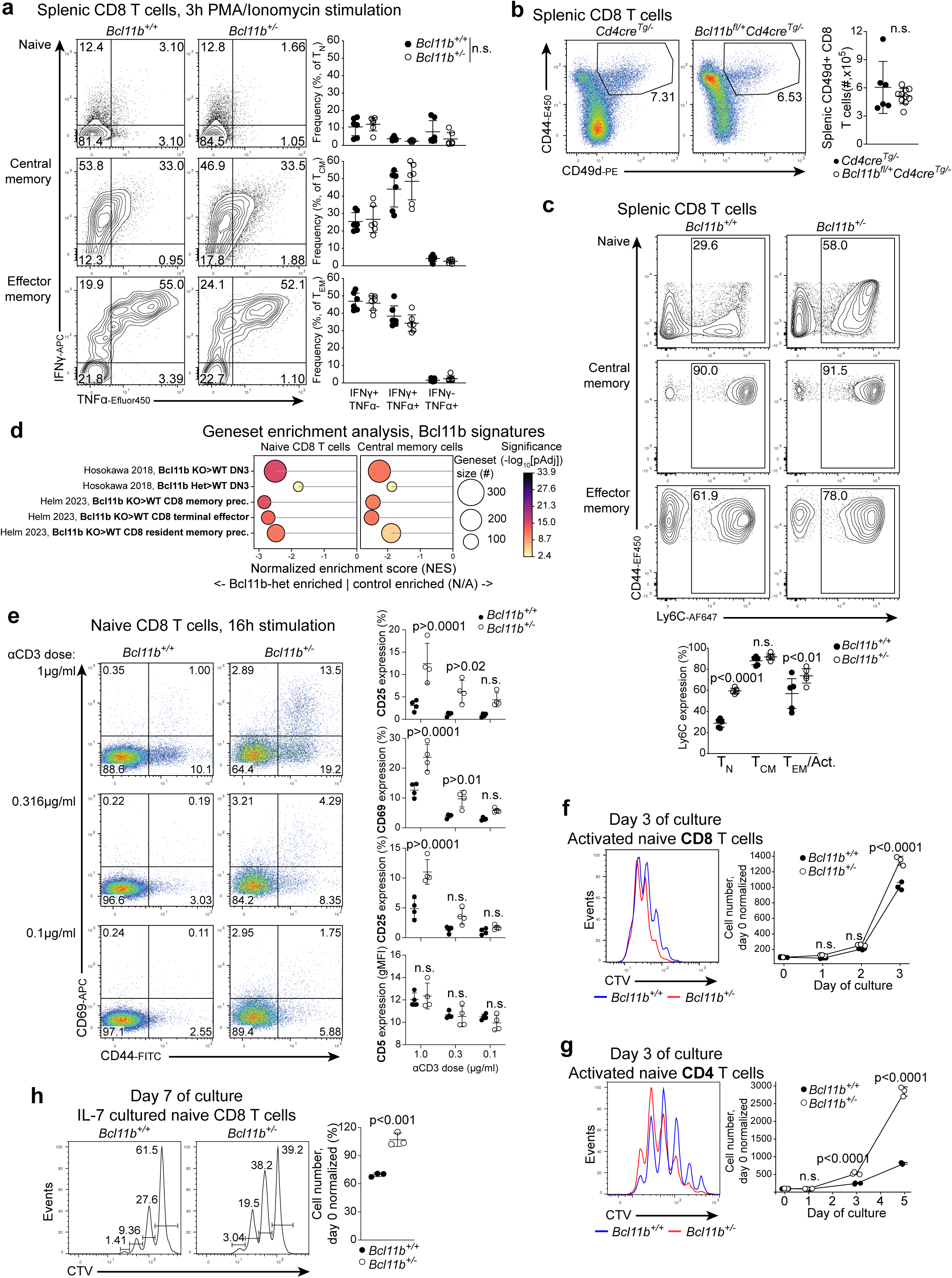
**a.** As in Figure 3c. Representative flow cytometry plots of IFN-γ and TNF-α expression of T_N_, T_CM_ and T_EM_ CD8 T cell subsets of from *Bcl11b^+/-^* and control spleens. **b.** Representative flow cytometry plots (left) and quantification (right) of CD49d+ ‘true’ memory T cells among *Bcl11b^fl/+^Cd4-cre^Tg^* and Cre-control splenic CD8T cells. **c.** Representative flow cytometry plots of the distribution of CD44 and Ly6C expression among T_N_, T_CM_ and T_EM_ CD8 T cell subsets of from *Bcl11b^+/-^* and control spleens, and quantification (lower) of Ly6C expression. **d.** As in Figure 3g. Geneset enrichment analysis of signatures of Bcl11b deleted populations from published datasets, comparing *Bcl11b^+/-^* and control T_N_ and T_CM_ populations. **e.** Naïve *Bcl11b^+/-^* and control CD8 T cells were cultured overnight in the indicated concentrations of plate-bound anti-CD3. Representative flow cytometry plots of CD69 and CD44 expression (left), and quantification of CD69, CD44, CD25 and CD5 expression (right) after culture. Naïve *Bcl11b^+/-^* and control CD8 (**f.**) and CD4 (**g.**) T cells cultured in anti-CD3 and IL-2 for three days and proliferation assessed by flow cytometry. Representative histograms of division tracing dye dilution (left), and quantification of cell numbers (right). **h.** Naïve *Bcl11b^+/-^* and control CD8 T cells cultured in IL-7 for a week. Left, representative histograms of dilution of division tracing dye, right, quantification of cell numbers after culture. Significance tested using one-way ANOVA with Šidák’s correction for multiple comparisons (a, c, e), two-way ANOVA with Tukey’s test for multiple comparisons (f, g), or Student’s two-tailed t test (b, h).

**Figure 4 – figure supplement 1.**
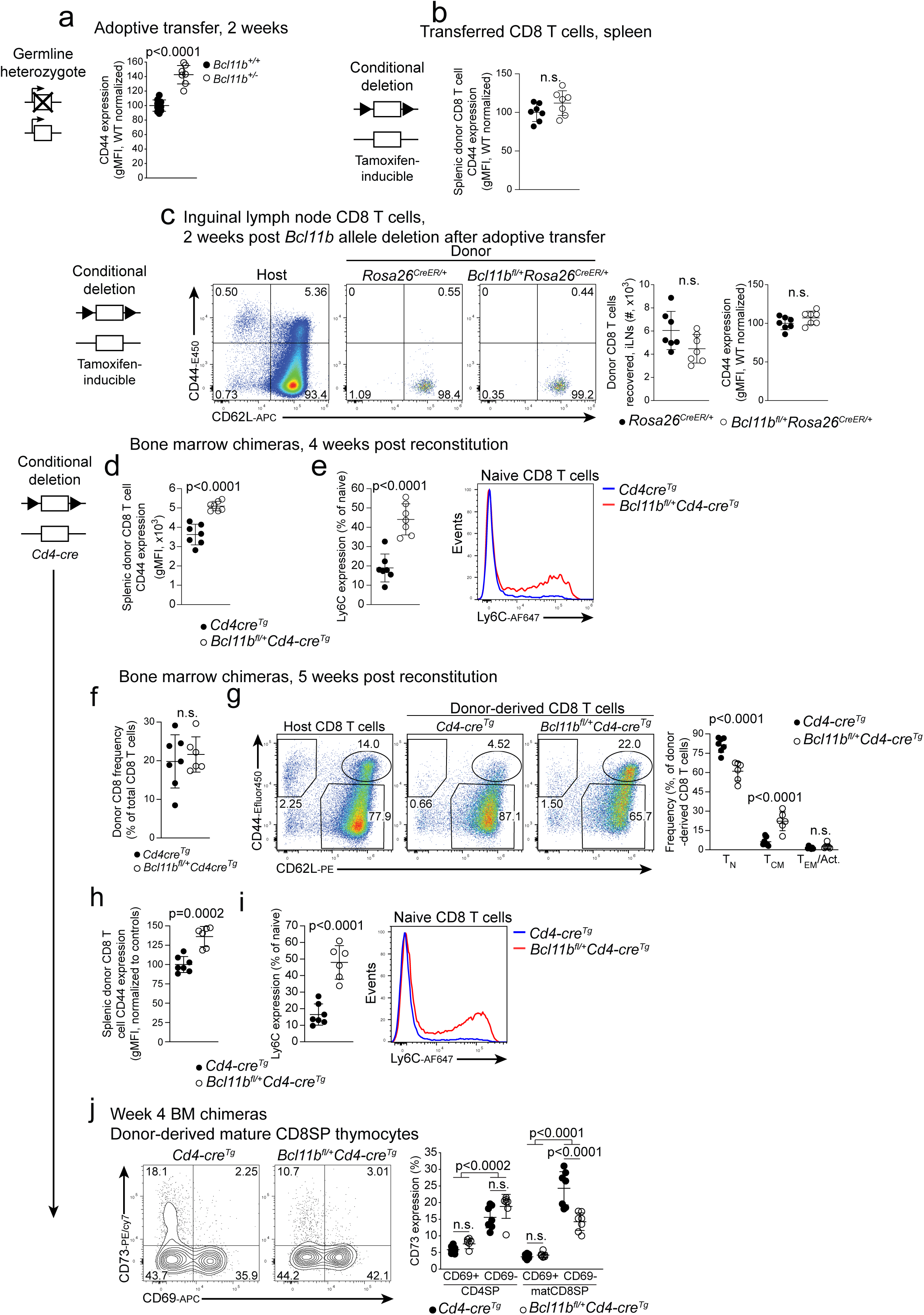
**a.** Quantification of CD44 expression by donor-origin naïve CD8 T cells in Figure 4b. **b.** Quantification of CD44 expression by recovered donor-origin CD8 T cells in Figure 4c. **c.** As in Figure 4c, for the cells recovered from pooled brachial, axial and inguinal lymph nodes. **d.-j.** Bone marrow chimeras generated from *Bcl11^fl/+^Cd4-cre^Tg^* and Cre-expressing donors analysed at the indicated timepoints post transfer. **d.** CD44 expression by recovered naïve donor-origin cells at 4 weeks. **e.** Quantification (left) and representative histograms (right) of Ly6C expression by donor-origin naïve CD8 T cells at 4 weeks. **f.-g.** As for Figure 4e, at 5 weeks post reconstitution. **h.** and **i.**, as for **d.** and **e.**, at 5 weeks post reconstitution. **j.** Representative flow cytometry plots (left) and quantification (right) of CD73 expression by mature single positive thymocytes 4 weeks post reconstitution. Significance tested using one-way ANOVA with Šidák’s correction for multiple comparisons (g, j) or Student’s two-tailed t test (a-f, h, i).

**Figure 5 Figure supplement 1.**
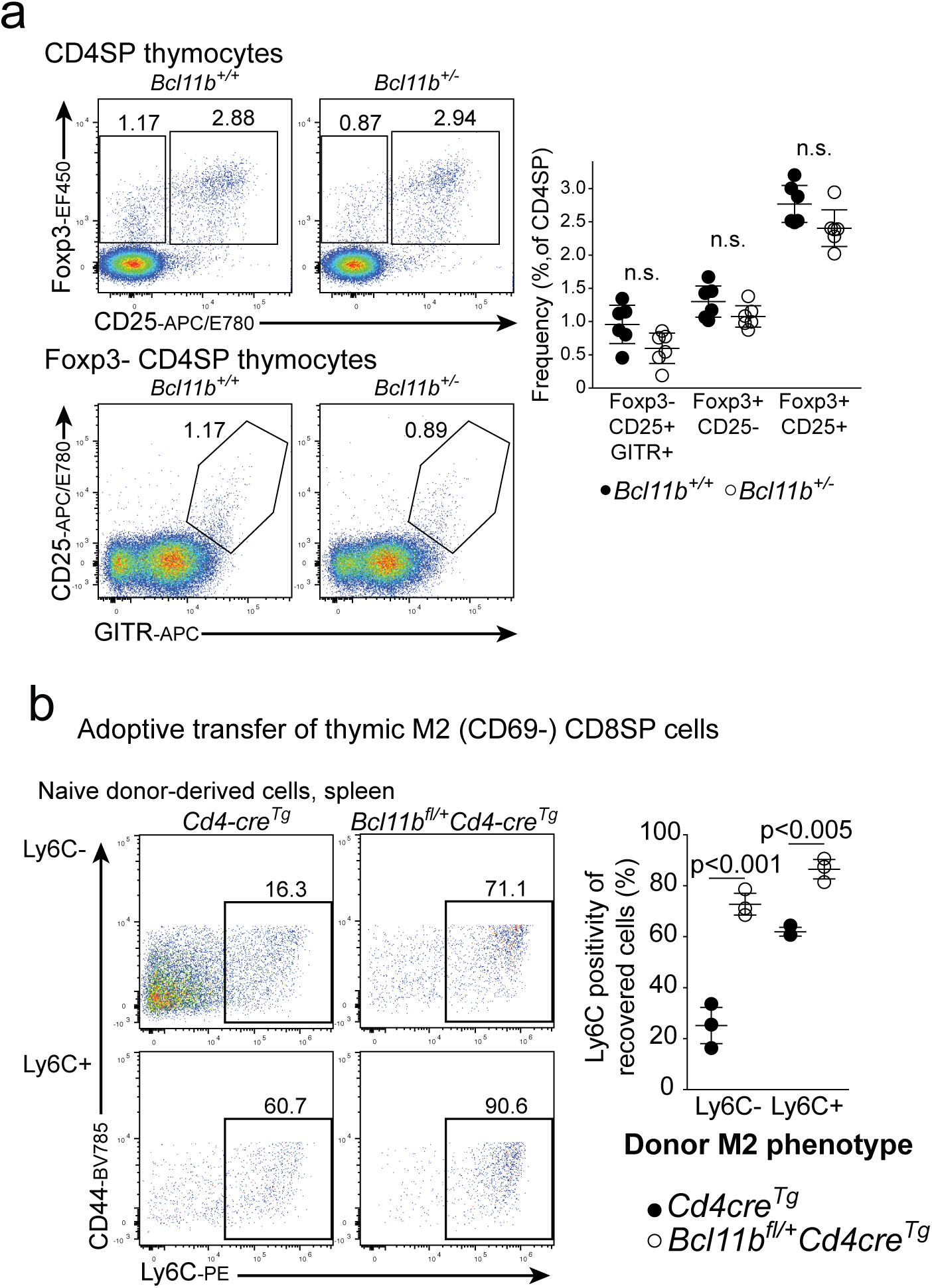
**a.** Representative flow cytometry plots (left) and frequencies (right) of thymic regulatory T cell lineage cells among CD4 single positive thymocytes from *Bcl11b^+/-^* and littermate control animals. **b.** Representative flow cytometry plots (left) and quantification (right) of Ly6C expression by recovered naïve CD8 T cells in Figure 5d. Significance tested using one-way ANOVA with Šidák’s correction for multiple comparisons

**Figure 6 – figure supplement 1.**
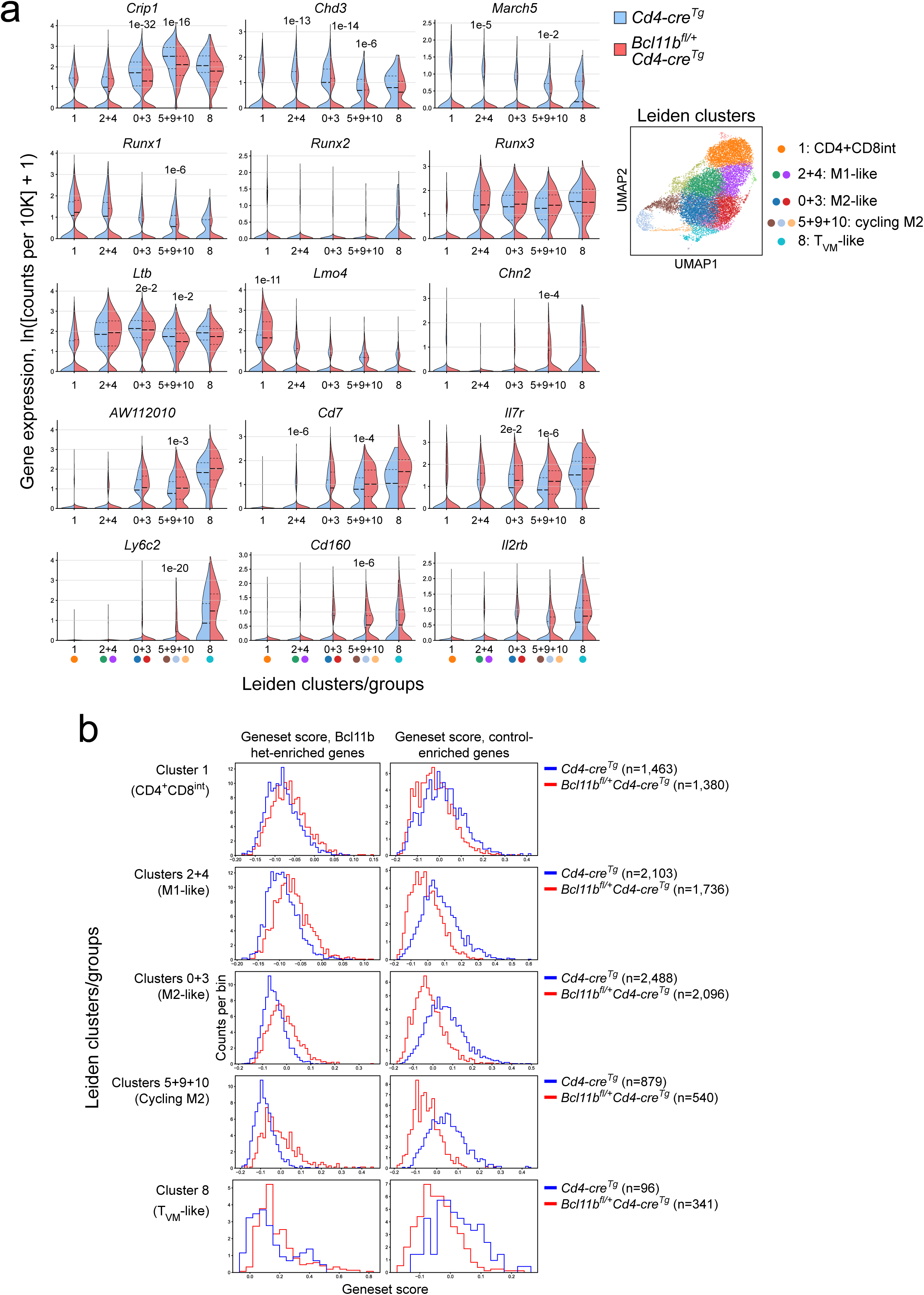
Single cell RNA-seq of thymic CD4^+^CD8^int^ and mature CD8SP cells, as in Figure 6. **a.** Violin plots of expression values of hypothesis-relevant and control genes, within indicated clusters or pooled clusters. Note that the small cell number in T_VM_-like cluster 8 penalized the statistical significance of comparisons. **b.** Histogram of geneset score values for genes ever found significantly (pAdj < 0.05, fold change > 2) enriched in either control or conditionally Bcl1b haploinsufficient cells in any cluster or pool of clusters by pseudobulk differential expression calling. Significance tested by DESeq2 (a).

**Figure 6 – figure supplement 2.**
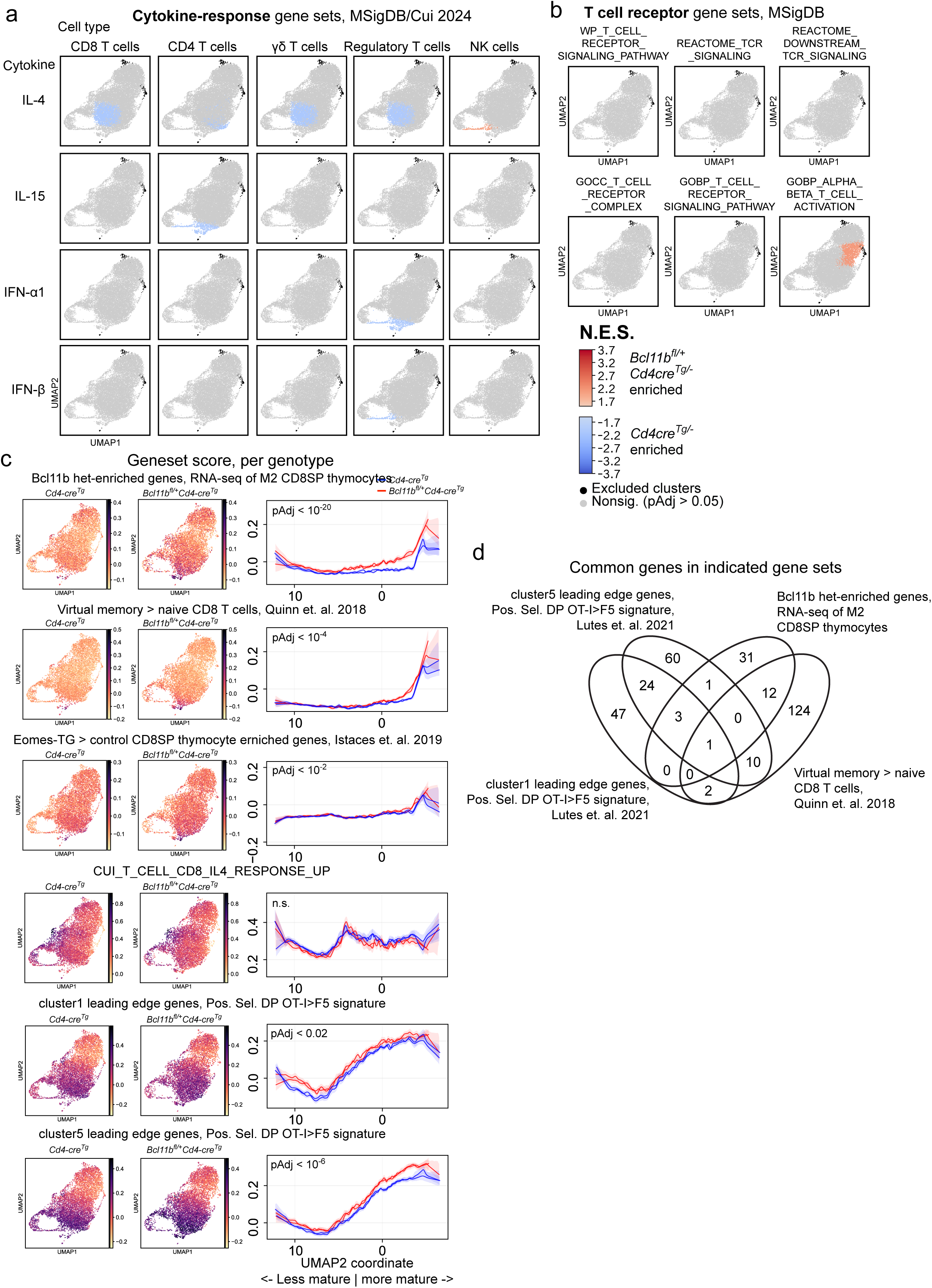
**a.-b.** Pseudobulk differential expression calls comparing genotypes in each cluster were treated to geneset enrichment analysis. Clusters are colored by normalized enrichment score (N.E.S.), with non-significant clusters grey and clusters insufficiently populated for testing colored black. **a.** Genesets of lymphocyte responses to cytokine stimulation. **b.** MSigDB genesets of TCR signaling. **c.** Geneset score for indicated gene sets was calculated per genotype and per sample. Left: featureplots showing enrichment of indicated gene set scores for each genotype in UMAP space. Right: Plots of mean gene set score values across UMAP2 bins. **d.** Venn diagram indicating common genes in the first four gene sets of panel c. Significance tested using a likelihood-ratio test comparing mixed-effects models of per-sample UMAP2-bin mean scores, with sample as a random intercept; the full model included a genotype x UMAP2-bin interaction and was compared to a reduced additive genotype + UMAP2-bin model (c),

**Figure 7 – figure supplement 1.**
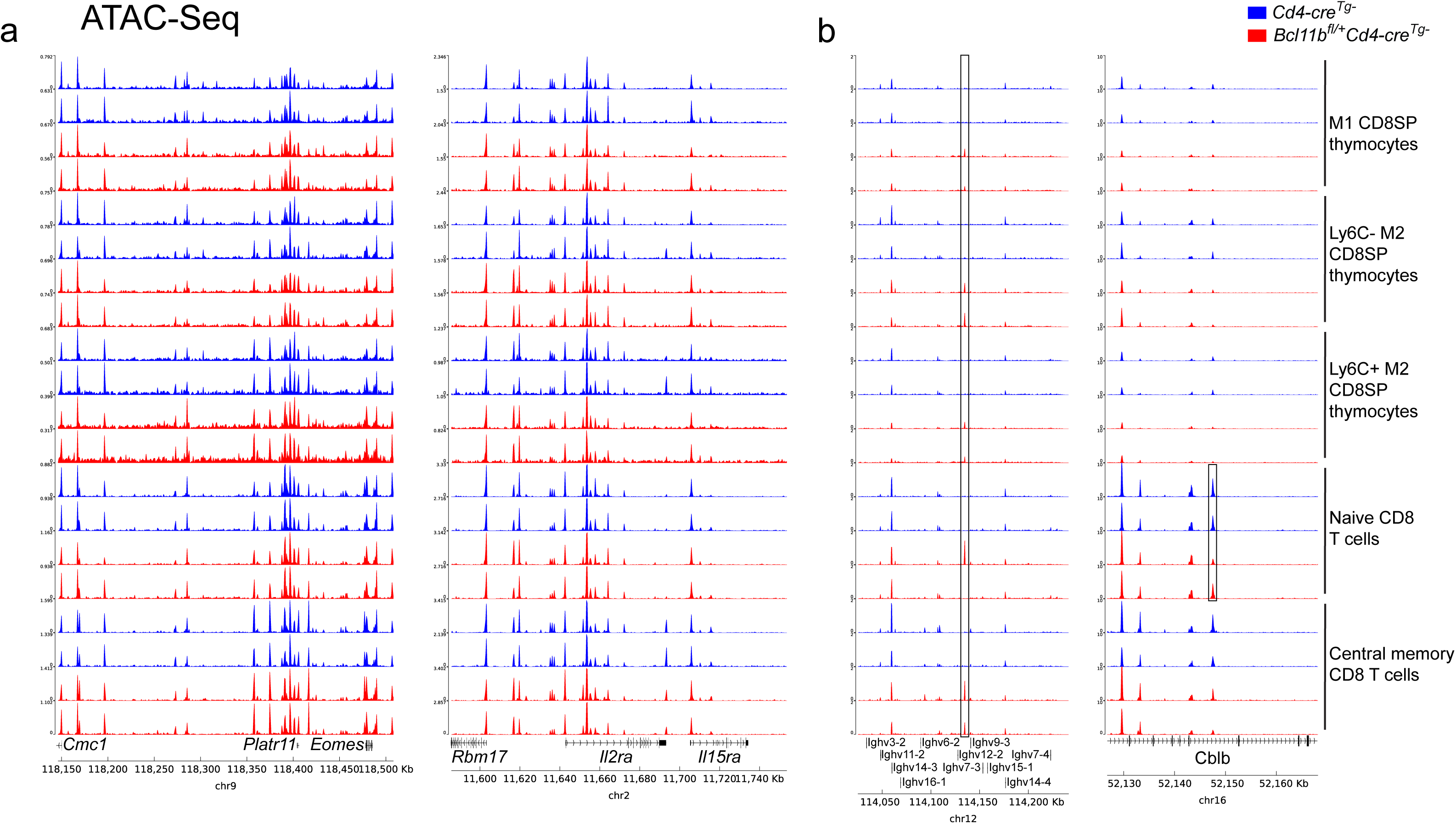
Representative genome browser tracks for all ATAC-seq replicates for indicated loci, CPM normalized. **a.** T cell memory relevant loci. **b.** Loci containing open chromatin regions statistically significantly (pAdj < 0.05) differentially accessible between genotypes (specific OCRs indicated by black boxes).

**Figure 8 – figure supplement 1.**
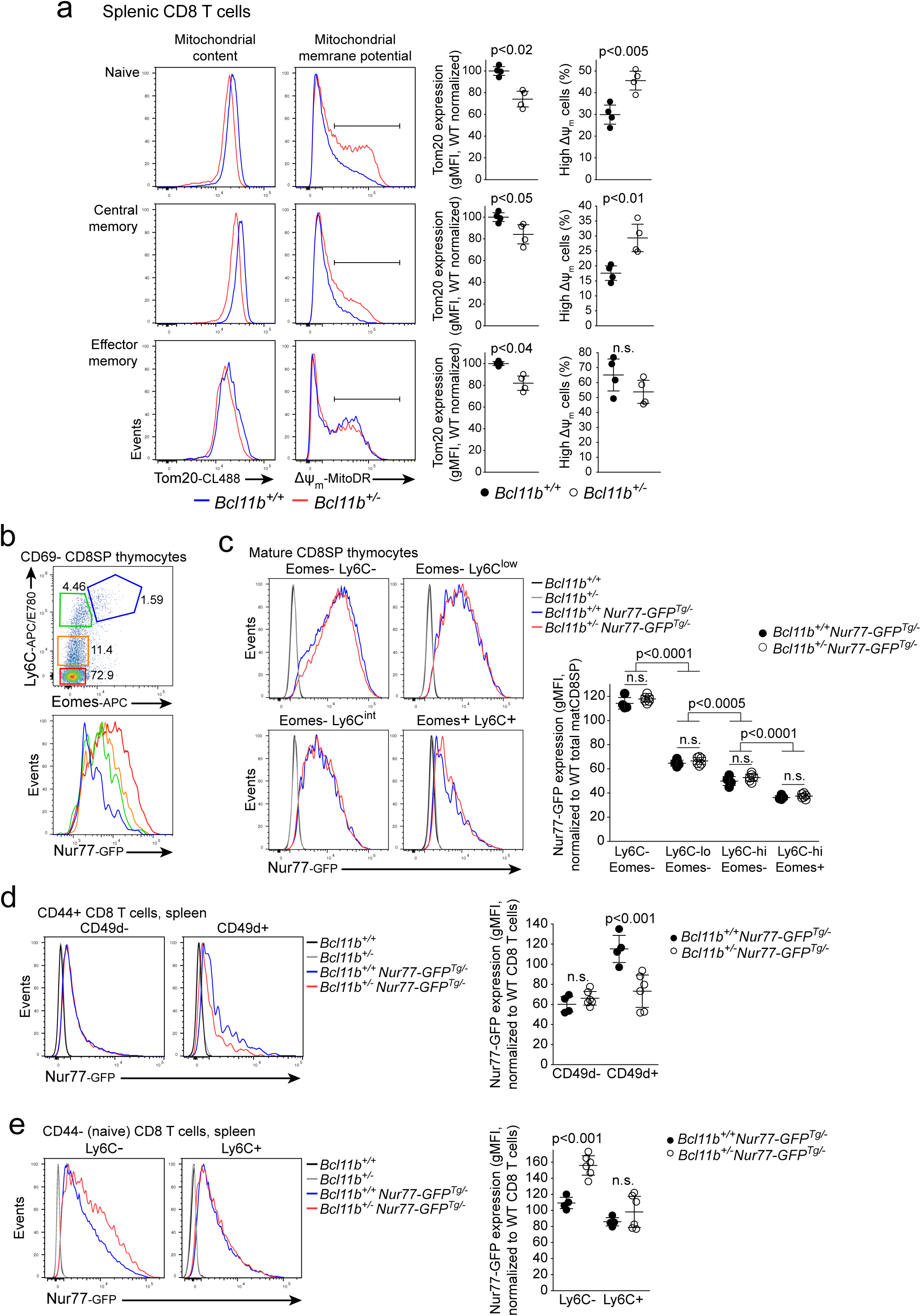
**a.** Flow cytometric analysis of mitochondrial content and polarization in splenic CD8 T cells. Representative histograms of Tom20 (left) and MitoTracker DeepRed FM (right) straining among *Bcl11b^+/-^* and littermate control spleens, quantified below. **b - e.** Flow cytometric analysis of Nur77-GFP reporter expression by *Bcl11b^+/-^* and littermate controls. **b.** Representative flow cytometry plot (upper) illustrating polygon gates used for panel c (upper) and representative histograms (lower) of Nur77-GFP expression within each. **c.** Representative histograms (left) of Nur77-GFP expression by *Bcl11b* heterozygote or control Ly6C and Eomes expressing CD8SP populations, quantified right. **d.** Representative histograms of Nur77-GFP expression among CD49d-negative T_VM_ and CD49d-positive ‘true’ memory cells of each genotype (left), quantified right. **e.** Representative histograms of Nur77-GFP expression among Ly6C-postive and negative naïve CD8 T cells of each genotype (left), quantified right. Significance tested using Student’s two-tailed t test (a), or one-way ANOVA with Tukey’s (c) or Šidák’s (d, e) correction for multiple comparisons.

**Supplementary table 1**

Differential expression analysis and normalized expression tables of Bcl11b heterozygous or homozygous loss in cultured DN3-like cells from Hosokawa et. al. 2018

**Supplementary table 2**

Differential expression analysis and normalized expression tables of Bcl11b heterozygous and control naive CD8 T cell populations from bulk RNA-seq.

**Supplementary table 3**

Differential expression analysis and normalized expression tables of Bcl11b heterozygous and control central memory CD8 T cell populations from bulk RNA-seq.

**Supplementary table 4**

Differential expression analysis and normalized expression tables of Bcl11b heterozygous and control M2 CD8SP thymocyte populations from bulk RNA-seq

**Supplementary table 5**

Differential accessibility analysis Bcl11b heterozygous and control thymic and splenic populations from bulk ATAC-seq. Reports fold changes and adjusted p values between genotypes for M1, Ly6C- M2 and Ly6C+ M2 thymocytes, and Naïve and central memory CD8 T cells from the spleen and lymph nodes.

**Supplementary table 6**

Pseudobulk analysis of Bcl11b haploinsufficient cells, controlled by cluster. DESeq2 output for genotype comparisons of indicated clusters/groups of clusters, list of significantly ‘WT’ and ‘Het’ genes pooled across comparisons, and gene lists from the Venn diagram in July_Figure 6 – figure supplement 2d.

